# THERM-D Uncovers Distinct Neural Mechanisms Separating Morning and Evening Body Temperature Rhythms in *Drosophila*

**DOI:** 10.64898/2026.02.19.706827

**Authors:** Tadahiro Goda, Olivia M. Lopez, Richard Ramolete, Nils Reinhard, Nick Ulle, Ayumi Fukuda, Yujiro Umezaki, Kazim Rizvi, Madison G Aikawa, Roslyn Catiis, Victoria Z. Marquez, Richard Ngo, Nikita Raj, Cora Fisher, Gregory T. Bui, Jalen Lee, Charlotte Helfrich-Förster, Taishi Yoshii, Fumika N. Hamada

**Affiliations:** Department of Neurobiology, Physiology and Behavior, University of California, Davis, Davis, CA 95616; Neurobiology and Genetics, Biocenter, University of Würzburg, Am Hubland, 97074 Würzburg, Germany; DataLab: Data Science and Informatics, University of California Davis, One Shields Avenue, Davis CA 95616; Graduate School of Environmental, Life, Natural Science and Technology, Okayama University, Okayama 700-8530, Japan

**Keywords:** circadian clock, body temperature rhythms, CRY, CRY-negative clock neurons and *Drosophila melanogaster*

## Abstract

Animal body temperature rises throughout the day and peaks in the evening, a pattern conserved across diurnal endotherms and ectotherms. However, the mechanisms driving the robust body temperature rhythms (BTR) remain largely unclear. Here, we developed a machine learning–based platform, temperature homeostasis evaluation of rhythmicity in model *Drosophila* (THERM-D), enabling continuous, high-throughput analyses of BTR. Using THERM-D, we identified robust BTR patterns reflecting flies’ morning and evening behaviors and revealed the function of CRYPTOCHROME (CRY)-negative clock neurons. About half of all clock neurons lack CRY, yet their function was unclear. Newly developed Gal4 drivers targeting CRY-negative neurons demonstrated that these neurons control the morning temperature rise without affecting evening BTR. The data suggests that separate clock circuits regulate morning and evening BTR. Thus, THERM-D elucidated the role of CRY-negative clock neurons, which are specialized for BTR regulation and distinct from the circuits controlling sleep-wake cycles.

## INTRODUCTION

Body temperature rhythm (BTR) is one of the most robust circadian rhythms and plays a crucial role in regulating metabolism and sleep (*1–4*). Body temperature drops during sleep and rises just before waking. This anticipatory response prepares animals for essential activities such as hunting and foraging. After this initial rise, body temperature continues to increase throughout the day and peaks before sleep onset (*4–11*). However, the mechanisms driving this rise and peak in body temperature remain largely unclear.

*Drosophila melanogaster* serves as a versatile model organism, offering powerful genetic tools that facilitate the mapping of neural circuits and the dissection of molecular mechanisms. We previously demonstrated that flies exhibit a robust temperature preference behavior (*12*). Through repeated behavioral assays at eight discrete time points throughout the day, we discovered that flies consistently prefer cooler temperatures in the morning and warmer temperatures in the evening, revealing a temperature preference rhythm (TPR) that is regulated by the circadian clock (*13*). Because flies are small ectotherms, their body temperature closely reflects the ambient temperature, suggesting that their TPR is effectively equivalent to their BTR (*14–16*). Using the TPR assays, we demonstrated that *Drosophila* BTR shares fundamental molecular mechanisms with those of mammals (*17, 18*). Both *Drosophila* and mammals exhibit a typical pattern of BTR: body temperature increases robustly in the morning and peaks in the evening.

However, several technical barriers within TPR assays rendered consistent BTR monitoring difficult (see Discussion). Since the TPR assay lasted only 30 minutes due to the lack of food, each assay required 40 repetitions to create a full 24-hour curve. A major concern was that the flies were woken up abruptly during the night because they were introduced to the apparatus across a full 24-hour day. This fact could not be ignored because of the close relationship between sleep initiation and body temperature decrease (*19, 20*). Therefore, a new behavioral assay was needed to understand the mechanisms by which temperature increases in the morning and peaks in the evening. This assay would continuously monitor the flies’ BTR with higher resolution, avoid disrupting their natural sleep-wake cycles, and detect more detailed behavior changes.

Here, we developed a machine learning-based analytical pipeline named THERM-D (<u>T</u>emperature <u>H</u>omeostasis <u>E</u>valuation of <u>R</u>hythmicity in <u>M</u>odel *<u>D</u>rosophila*) that constantly monitors fly BTR and processes data more efficiently than the TPR assay. THERM-D enables continuous and accurate BTR data for over 24 hours to obtain intrinsic BTR, as the flies were fed and not disturbed in their sleep-wake cycles. Using THERM-D, we demonstrate that *Drosophila* exhibit a robust BTR over several days, revealing more detailed and physiologically relevant behavioral changes.

The blue-light sensor CRYPTOCHROME (CRY) plays an important role in regulating circadian rhythms in *Drosophila* (*21, 22*). CRY-positive clock neurons mainly regulate sleep-wake cycles via light entrainment. However, CRY is not expressed in about half of all clock neurons (*23, 24*) and the function of these CRY-<u>negative</u> clock neurons have remained largely unclear. Recent studies indicate that CRY-negative clock neurons are more prone to temperature entrainment (*25*) and that several CRY-negative clock neurons interact with temperature-processing neural circuits (*26–28*). These findings suggest that CRY-negative clock neurons may play a key role in regulating BTR. However, the lack of genetic tools that specifically target each CRY-negative clock neuron has hindered the ability to evaluate their role directly.

Here, we identify several GAL4 drivers that selectively label CRY-negative clock neurons and demonstrate that these neurons function as novel oscillators controlling BTR in the morning. This finding highlights a mechanistic distinction between the regulation of morning and evening BTR. This result further supports the idea that BTR is regulated by neural circuits different from of those regulating the sleep–wake cycle since CRY-negative clock neurons do not play a major role in sleep-wake regulation. Altogether, our machine learning-based THERM-D assay provides a powerful new platform for dissecting the neural and molecular mechanisms underlying the coordination between circadian timekeeping and physiological temperature homeostasis.

### HIGHLIGHTS

1. THERM-D, a novel machine-learning-based assay, enables continuous monitoring of body temperature rhythms (BTR) in *Drosophila* for over 24 hours.
2. CRY-negative clock neurons play a critical role in driving BTR in the morning.
3. Distinct mechanisms regulate BTR in the morning and evening.
4. BTR and sleep–wake cycles are controlled by different neural circuits.

## RESULTS

### THERM-D: A novel machine learning-based BTR assay in *Drosophila*

*Drosophila* exhibit robust temperature preference behavior (*12*). In the process of repeating the temperature preference behavioral assays, we discovered that flies exhibit temperature preference rhythm (TPR), preferring a lower temperature in the morning and a higher temperature in the evening (Fig. S1A). Due to their small size as ectotherms, the fly’s body temperature is about the same as their surrounding temperature and therefore, TPR indicates their BTR (*13, 14*). Using the TPR assays, we demonstrated that *Drosophila* BTR and mammalian BTR share fundamental molecular mechanisms (*17, 18*). However, the TPR assay has several disadvantages, particularly the disturbance of fly sleep-wake cycles at the time they are applied to the apparatus in the night (Fig. S1A) (see Discussion). To overcome this limitation, we developed a new method, THERM-D, which enables continuous monitoring without disturbance of the flies’ sleep-wake cycles and permits efficient data analysis (Fig. 1A). With the THERM-D assay, data analysis requires examining about 42 photos. To expedite data acquisition of the fly’s location and calculation of the fly’s body temperature, we developed a machine learning algorithm (see the Supplemental Information for details). Thanks to this algorithm, we can now analyze the data in just a few minutes, saving a significant amount of time. Previously, TPR assays took several months to investigate one genotype to generate a 24-hour curve.

**Figure 1:**
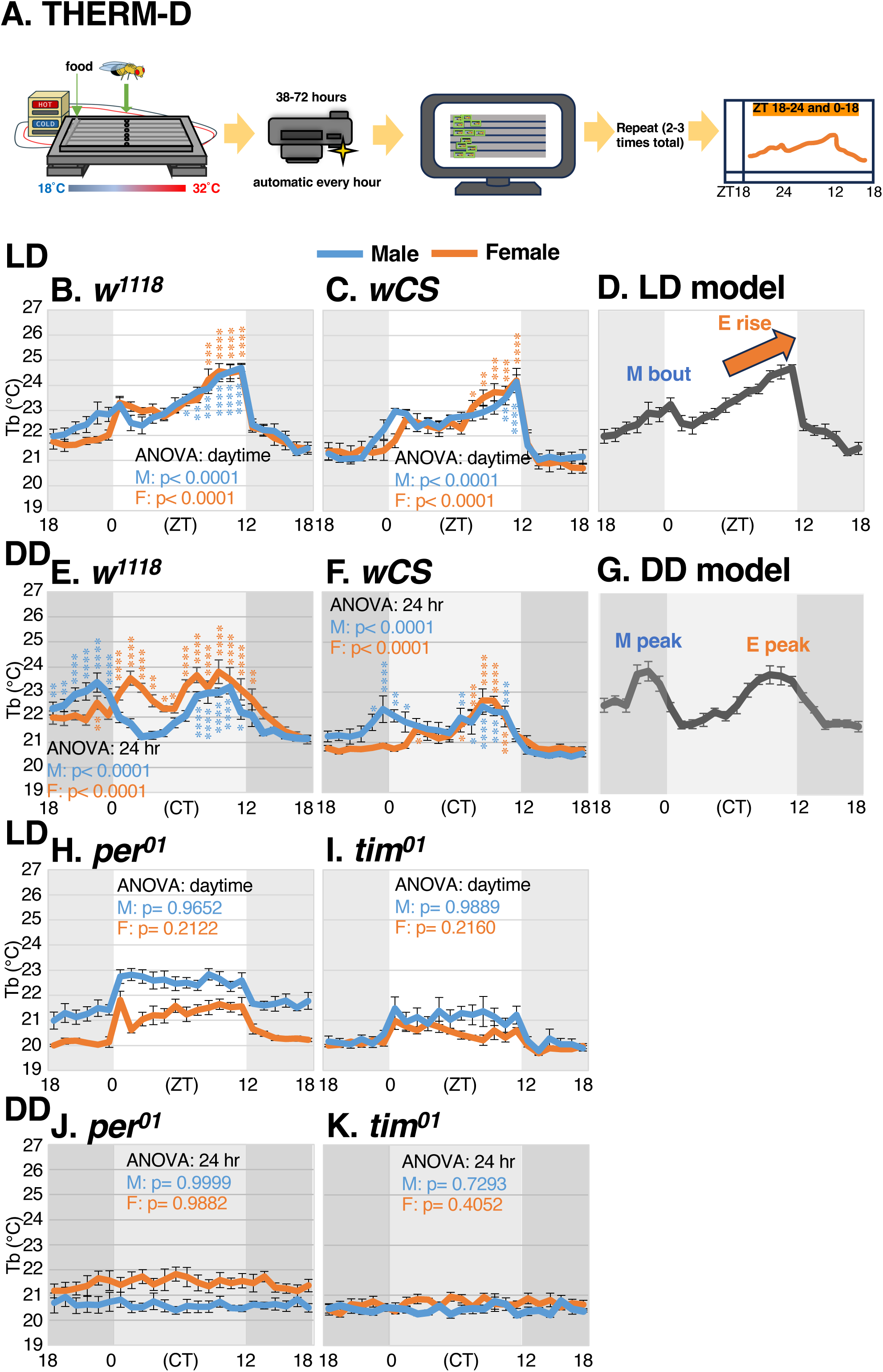
Machine learning-based BTR assay THERM-D show innate BTR. **(A)** THERM-D assay used to monitor *Drosophila* BTR. Flies are inserted into a 6-lane plexiglass chamber atop a metal apparatus with Peltier devices that maintain a temperature gradient of 18 to 32°C. Fly body temperature rhythm (BTR) is continuously monitored and recorded each hour for 2 to 3 days. Subsequent data is analyzed via machine learning platforms. Assays were performed to a total of n = 6 ∼ 9. **(B, C)** *w^1118^* (B) and *wCS* (C) tested using THERM-D assays in LD. The male and female flies were tested separately and are shown in the graphs by blue and orange lines, respectively. **(D)** LD model of *Drosophila* BTR characterized from male *w^1118^*; morning (M) bout occurs ZT 18-6 (18-24 through 0-6), E rise occurs ZT 6-12. One-way ANOVA and Dunnett’s multiple comparisons test with the lowest body temperature (Tb) in ZT 0-12 were used to analyze E rise. *p < 0.05. **p < 0.01. ***p < 0.001. ****p<0.0001. **(E, F)** *w^1118^* (E) and *wCS* (F), tested using THERM-D assays in DD. **(G)** DD model of *Drosophila* BTR characterized from male *w^1118^* flies showing M peak and E peak. One-way ANOVA and post hoc Dunnett’s multiple comparisons test with the lowest temperature point of the BTR curve were used for M and E peaks in DD. *p < 0.05. **p < 0.01. ***p < 0.001. ****p<0.0001. **(H, I)** BTR phenotype of clock knockout mutants, *per^01^* (H) and *tim^01^* (I), in LD. **(J, K)** BTR phenotypes of clock knockout mutants, *per^01^* (J) and *tim^01^* (K), in DD. Dark shadowed regions on the graphs represent night; light regions represent day; medium shadowed regions represent subjective day in DD.

### The innate BTR behavior

To examine BTR behavior using THERM-D, we tested wild-type-like (WT) male and female *w^1118^* flies separately. Under light-dark (LD) conditions, both sexes exhibited significant BTR; body temperature initially rises in the morning, dips slightly during midday, and then climbs again until reaching a prominent peak in the evening (Fig. 1B). The one-way ANOVA during the daytime (ZT 0-12) showed that the body temperature of *w^1118^* males at ZT 6-12 was significantly higher compared to the lowest point of daytime body temperature at ZT 2. Another WT fly strain, white *Canton-S* (*wCS*), which has a CS background with the *w^1118^* mutation, exhibited a similar pattern to *w^1118^* (Fig. 1C). Thus, we named the first phase of the temperature rise during ZT 18-6 (18-24 through 0-6) as the morning bout (M bout) and the rise during ZT 6-12 as the evening rise (E rise) (Fig. 1D). Notably, we observed sexual dimorphism within BTR. The *w^1118^* and *wCS* males began raising their body temperature before the start of the daytime (ZT 0), but females did not (Fig. 1B and C). This initiation of body temperature rise in males was significant as calculated by Simple Linear Regression (SLR) (*w^1118^*: p = 0.0071, *wCS*: p < 0.0001).

In constant darkness (DD), the *w^1118^* and *wCS* flies exhibited robust fluctuations with significant double peaks occurring in the morning and evening (Fig. 1E and F), named as the morning peak (M peak) and evening peak (E peak) (Fig. 1G). In these flies, males and females both displayed a robust M peak; however, the M peak of females occurred three hours after the M peak of males (Fig. 1E and F). Thus, in females, the rise in body temperature begins after the start of the subjective day (CT 0) in DD (Fig. 1E and F) and this delayed phenotype is also evident during the daytime in LD (Fig. 1B and C). On the other hand, the E peaks occurred at the same time for both sexes, though they both showed a phase advance of a few hours compared to the LD phenotype (Fig. 1B, C, E and F).

Notably, the rhythms of BTR in DD persisted for several days, but the amplitude was strongest during the first day and dampened after second day (Fig. S1B). The quality of food, deteriorating and drying out over time, is likely affecting the behavior since the internal metabolic state strongly affects body temperature (*29*). Additionally, in DD, each individual fly’s circadian clock runs at its own intrinsic period. Because we are monitoring the average body temperature of approximately 10–15 flies, each fly’s rhythm is likely to become out of phase over time. Therefore, we focused on the interval from ZT (or CT)18-24 through 0-18, where the BTR is most robust in both LD and DD conditions, to investigate the mechanisms underlying BTR.

### BTR is clock dependent

To determine whether BTR is dependent on the circadian clock, we tested mutants of the circadian clock genes *period* (*per*) and *timeless* (*tim*): *per^01^* and *tim^01^*. The *per^01^*and *tim^01^* flies, both male and female, did not increase their body temperature during the daytime in LD (Fig. 1H and I). The flies showed a masking effect when the light was ON or OFF (*30*): light ON caused a sudden increase in temperature at ZT 0, and light OFF caused a sudden drop in temperature at ZT 12 (*13*). Therefore, light induces an increase in body temperature irrespective of the circadian clock. We also observed a similar effect using TPR assays (*13, 31*). Without circadian control, animals are more likely to react to lights turning on and off. Furthermore, the *per^01^* and *tim^01^*flies showed no BTR fluctuation in DD (Fig. 1J and K). These results indicate that BTR is controlled by the circadian clock. Taken together, our newly developed platform, THERM-D, is a robust and reliable behavioral assay that enables the accurate and systematic investigation of innate BTR in *Drosophila*.

### DN2s and DN1as play an important role in the rhythmicity of BTR in DD

The fly brain contains approximately 240 clock neurons (*26*). Based on their cell body size and location in the brain, clock neurons are divided into groups of lateral <u>n</u>eurons (LNs) and <u>d</u>orsal <u>n</u>eurons (DNs) (*32*). Previously, we demonstrated that dorsal <u>n</u>eurons <u>2</u> (DN2s) and <u>a</u>nterior <u>d</u>orsal <u>n</u>eurons <u>1</u> (DN1as) are key core clock neurons for BTR by using TPR assays in LD (*13, 33*).

To reevaluate the roles of DN2s and DN1as in BTR using THERM-D, we disrupted the circadian molecular clock by employing clustered regularly interspaced short palindromic repeats (CRISPR)/CRISPR associated protein 9 (Cas9) genome editing technique. A *timeless (tim)*-CRISPR line was developed and used to eliminate the circadian clock in clock neurons (*33–36*). First, we used *Clk856-Gal4* to verify *tim*-CRISPR, knocking out *tim* in all clock neurons. In DD, TIM-depleted male and female flies lost these M and E peaks while control flies (*Gal4/+* and *UAS/+*) showed robust M peak and E peak (Fig. 2A and S2A). In LD, control flies demonstrated a significant E rise in body temperature, but neither male nor female flies with *tim* deletion in all clock neurons showed E rise (Fig. 2B and S2B). The data showed that *tim*-CRISPR successfully drives abnormal BTR phenotypes and confirmed that the circadian clock plays an essential role in BTR for both LD and DD.

**Figure 2:**
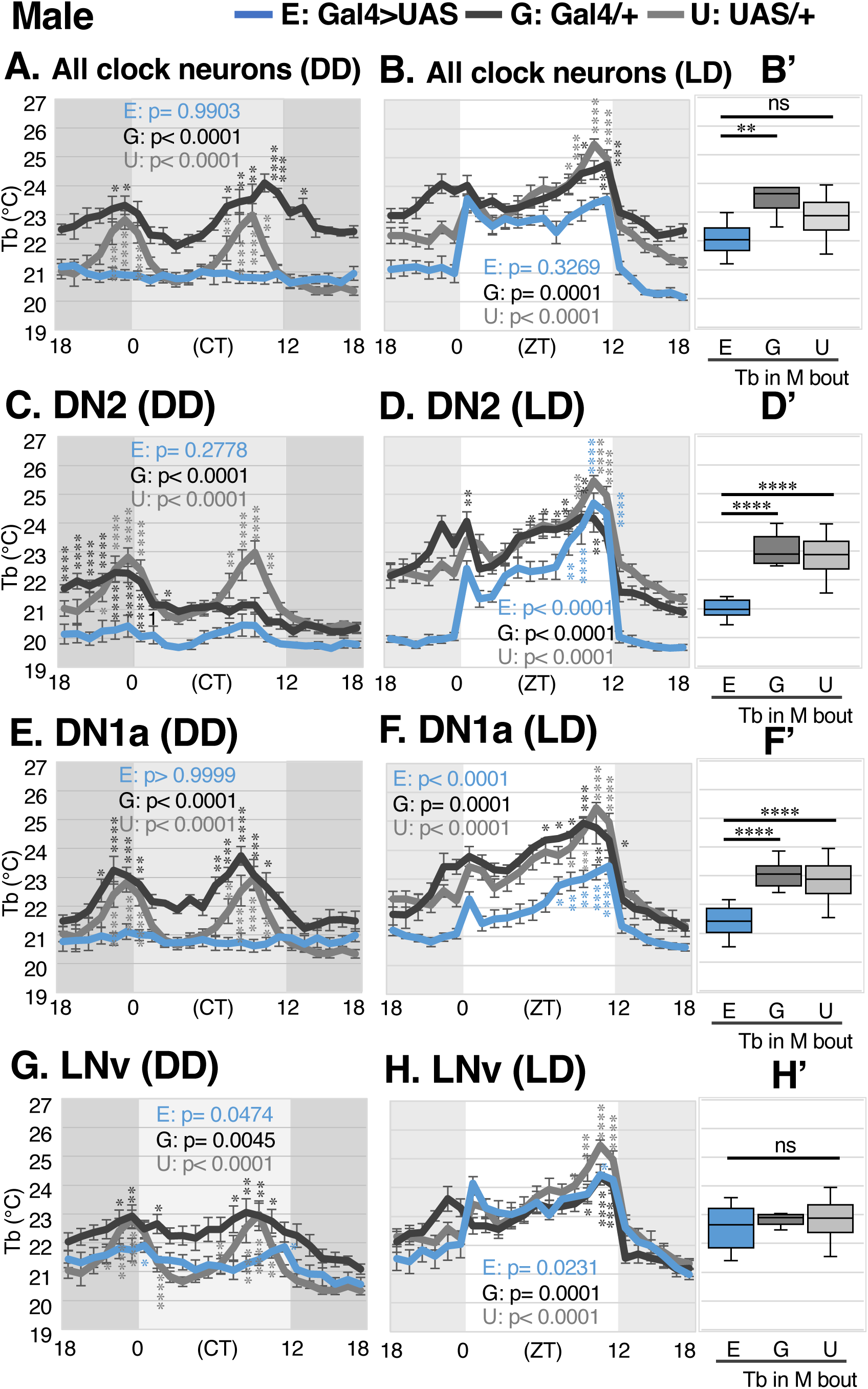
DN2 and DN1a control overall rhythmicity in DD and M bout in LD. **(A-H)** Comparison of BTR between clock-disrupted flies (blue) and control flies (*Gal4* (black) and *UAS* (gray)) in DD and LD. Clocks were disrupted via targeted *tim*-CRISPR expression in all clock neurons (A, B), DN2s (C, D), DN1as (E, F), and LNvs (G, H). One-way ANOVA and post hoc Dunnett’s test were used for comparison to the lowest body temperature (Tb) during the whole day (CT 18-24 and 0-18) in DD or daytime (ZT 0-12) in LD. **(B’, D’, F’, H’)** Comparison of average Tb between experimental flies (E), *Gal4* control (G), and *UAS* control (U) during M bout (ZT 18-24 and 0-6) in LD. ANOVA and post hoc Dunnett’s test were used for comparison between the experimental flies and *Gal4* control or *UAS* control. ∗∗∗∗p < 0.0001, ∗∗∗p < 0.001, ∗∗p < 0.01, or ∗p < 0.05 is shown.

Next, we reevaluated the roles of DN2s and DN1as for BTR using THERM-D in DD. In males, knocking out *tim* in either DN2s or DN1as resulted in significantly dampened M and E peaks (Fig. 2C and 2E). In particular, knocking out *tim* in DN2s caused an overall low body temperature throughout the day (Fig. S3F). In females, although knocking out *tim* in DN2s exhibited an abnormal phenotype with advanced E peak, they still sustained the rhythmicity (Fig. S2C). Furthermore, we found that knocking out *tim* in DN1as of female flies showed no BTR (Fig. S2E). Overall, the abnormal DD phenotype observed in TIM depletion in DN1as is generally stronger than that in DN2s. It is important to note that *DN2-Gal4* is only expressed in one of the two DN2s (*13*), while the *DN1a-Gal4* is expressed in both of the two DN1as. Therefore, the clock depletion in DN2 using *DN2-Gal4* did not completely block their functions, potentially leading to a partial abnormal phenotype. Together, our data suggest that clocks in DN2s and DN1as are important for BTR rhythmicity in DD.

### DN2s and DN1as play an important role in establishing body temperature setpoints M bout in LD

Next, we focused on M bout in LD. Notably, the timing of the rise in body temperature varies depending on the sex of the organism (Fig. 1B and C) and the *Gal4* and *UAS* genetic background (Fig. 2). Therefore, we compared body temperature between experimental and control flies to evaluate M bout (ZT 18-24 and 0-6).

Remarkably, body temperatures of males with clock-ablated DN2s and DN1as exhibited significantly lower body temperature in LD during M bout (Fig. 2D’ and 2F’). In females, TIM-depleted DN2s caused lower body temperatures only during ZT 18-24 but not during 0-6 (Fig. S2D”) whereas TIM-depleted DN1as caused lower body temperatures during the entirety of M bout (Fig. S2F’). These results demonstrate the importance of DN2 and DN1a clocks in the body temperature rise of M bout in LD. Unexpectedly, knocking out *tim* in DN2 or DN1a still showed a significant E rise in both males (Fig. 2D and 2F) and females (Fig. S2D and S2F) in LD. We thought that the roles of DN2s and DN1as may be redundant and knocking out TIM in either DN2s or DN1as might not be sufficient to cause an abnormal E rise. Therefore, to knock out *tim* in both DN2s and DN1as simultaneously, we employed two other Gal4s, *CrzR-Gal4* and *tim-Gal4; Pdf-Gal80.* While *CrzR-Gal4* drives expression in both DN1as, DN2s, and a few DN3s (Fig. S3E), *Tim-Gal4; Pdf-Gal80* drives expression in most dorsal clock neurons (*37*). The flies with *tim* knocked out using these Gal4 drivers resulted in significantly dampened M and E peaks in males in DD (Fig. S3A and S3C). However, the flies still showed no significant change in body temperature during M bout (Fig. S3B’ and S3D’, see discussion) and a significant E rise in LD (Fig. S3B and S3D). Therefore, we conclude that other clock neurons besides the DN2s and DN1as may contribute to E rise in BTR under LD conditions.

### LNvs are not the primary clock neurons responsible for BTR

LNvs are the core clock neurons of the sleep-wake cycle. Without a clock in the LNvs, flies do not exhibit sleep-wake cycles (*37–40*). In previous assays using TPR, we demonstrated that LNvs are not the primary clock neurons for TPR (*13, 41*). We reevaluated the role of LNvs using THERM-D. Knocking out *tim* in LNvs resulted in robust BTR in LD and DD conditions for both male and female flies (Fig. 2G, 2H, 2H’, S2G, S2H and S2H’). These results suggest that the clocks in LNvs are not the primary clock neurons responsible for BTR.

### LPNs are responsible for morning BTR: M bout in LD and M peak in DD

LPNs receive input from the warm-sensing neurons called Anterior Cells (AC) (*12*) and from high temperature-processing neurons (TPN-IV). Additionally, LPNs are involved in sleep (*42*) and temperature-dependent siesta sleep (*27*). Therefore, we sought to determine whether the LPN clock is involved in BTR regulation. In DD, flies with TIM-depleted LPNs exhibited a weak rhythm, but M and E peaks were not significant (Fig. 3A). The body temperature of the flies was significantly lower than that of the control group in DD (Fig. S3G). In addition, knocking out *tim* in LPNs resulted in significantly lower body temperatures during M bout (Fig. 3B’), but maintained a robust E rise (Fig. 3B) in LD. Female flies showed a substantially dampened M peak but robust E peak in DD (Fig. S4A) and significantly lower body temperatures during M bout with a robust E rise in LD (Fig. S4B and S4B’). The data suggest that LPNs play an important role in BTR regulation, especially in body temperature rise during M bout in LD and M peak in DD.

**Figure 3:**
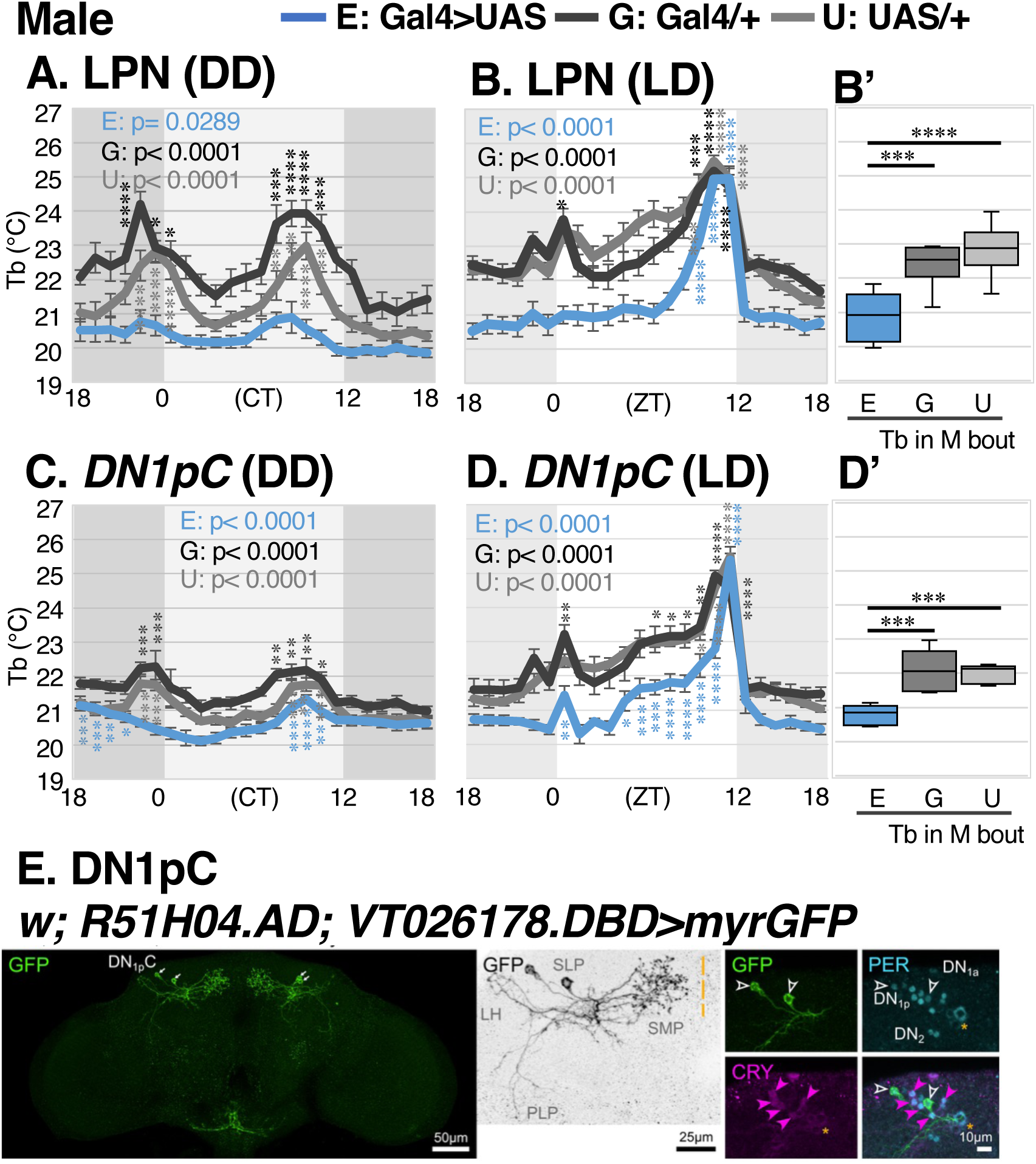
LPN and DN1pC control M peak in DD and M bout in LD. **(A-D)** Comparison of BTR between flies with clock disruption (blue) and control flies (*Gal4* (black) and *UAS* (gray)) in DD and LD. Clocks were disrupted via targeted *tim*-CRISPR expression in LPNs (A, B) and DN1pCs (C, D). **(B’, D’)** Average temperature comparison of M bout in LD between experiment (E), Gal4 control (G), and UAS control (U). Same statistical analysis as Fig. 2 was used. **(E)** Image of *spl-DN1pC-Gal4* (*w;R51H04.AD;VT026178.DBD*) crossed with *UAS-myrGFP*. *spl-DN1pC-Gal4* expressing neurons were labeled with GFP (green). Clock neurons were immunostained with anti-PERIOD (PER) antibody (cyan). CRY-positive neurons were immunostained with anti-CRY antibody (magenta). The yellow dashed line indicates the midline of the brain.

### DN1pCs also regulate morning BTR

Interestingly, like DN2s, LPNs do not express cryptochrome (CRY) (*25*). While CRY is an important blue light sensor that resets the molecular clock cycle in *Drosophila* (*21, 22*), several groups of clock neurons lack CRY expression. Among these CRY-negative oscillators, LPNs are known to receive temperature inputs (*27*). Therefore, we targeted a subset of the posterior dorsal clock neurons (DN1pC) which are also CRY-negative oscillators that receive temperature signals from antennal temperature-sensing neurons (ACs)(*26*). Because Gal4s that are specific to DN1pC were unavailable, we used flylight (*26*) and successfully identified Gal4 candidates that are selectively expressed in DN1pC (see material method). Using myrGFP to label neurons via immunostaining with PER and CRY antibodies, we revealed that the R51H04.AD;VT026178.DBD, *split-DN1pC-Gal4* driver is selectively expressed in DN1pC neurons (Fig. 3E).

To examine the role of DN1pC in BTR, we used this *split-DN1pC-Gal4* driver to selectively knock out *tim* in DN1pCs. In DD, both male and female flies with knocked out *tim* in DN1pCs lost their M peak but maintained a robust E peak (Fig. 3C and S4C). In LD, both male and female flies significantly lowered their body temperatures during M bout (Fig. 3D’ and S4D’) but maintained a robust E rise (Fig. 3D and S4D). The data suggest that DN1pC also plays an important role in the M bout of LD and the M peak in DD. Since DN2, LPN, and DN1pC are all CRY-negative neurons that regulate morning BTR, this suggests a potential pathway in which CRY-negative neurons regulate morning BTR.

### CRY-negative LNds and DN3s are involved in the regulation of morning BTR

We asked whether the other CRY-negative clock neurons, such as CRY-negative LNds and s-CPDN3As, might also regulate morning BTR. Because Gal4 tools expressed in these CRY-negative clock neurons were unavailable, we first identified two *Gal4s* that were selectively expressed in the CRY-negative LNds and s-CPDN3As, respectively (Fig. 4E and F) (*26*) (see material methods).

**Figure 4:**
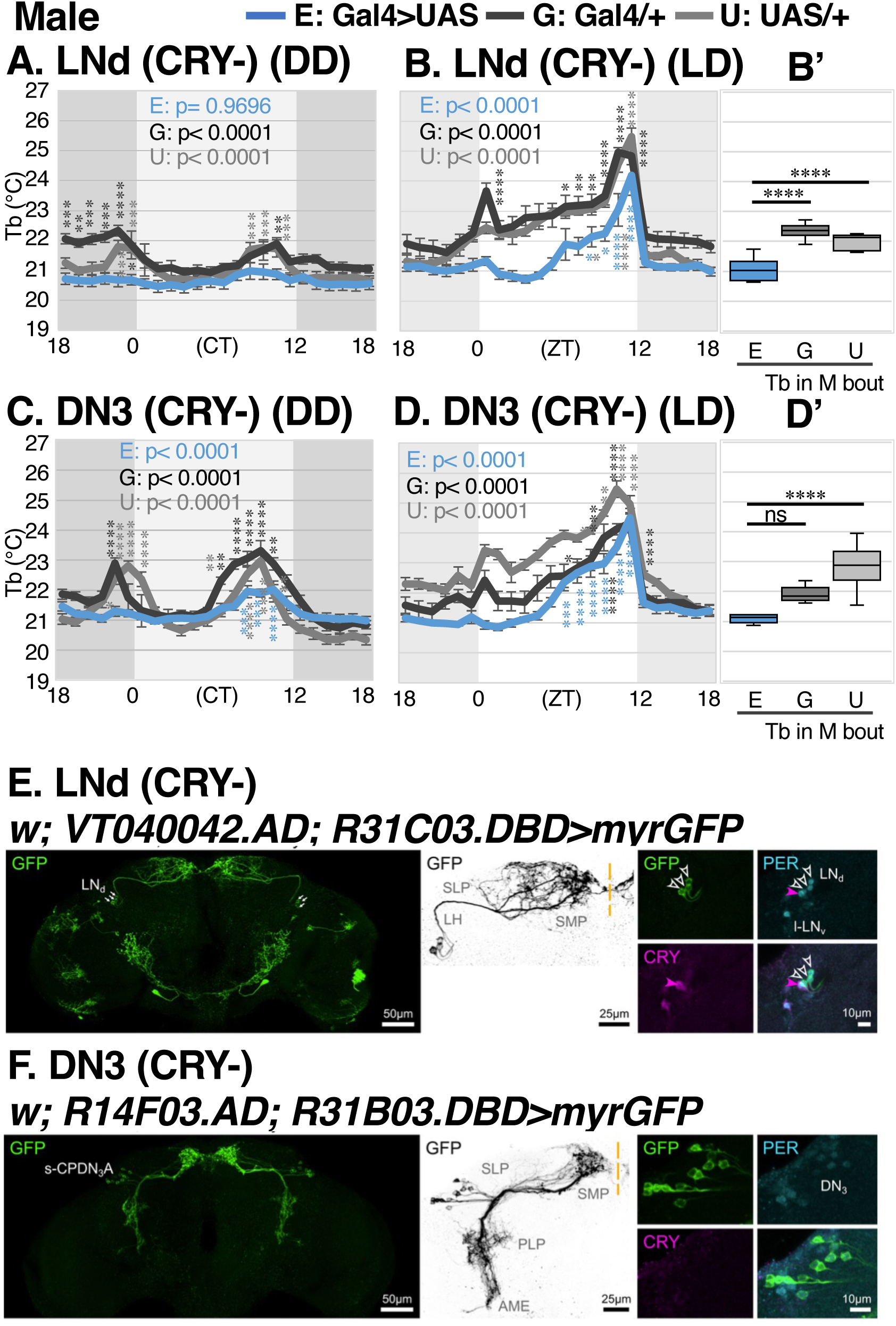
CRY-negative LNd and s-CPDN3A control M peak in DD and M bout in LD. (**A-D**) Comparison of BTR between flies with clock disruption (blue) and control flies (*Gal4* (black) and *UAS* (gray)) in DD and LD. Clocks were disrupted via targeted *tim*-CRISPR expression in CRY-negative LNds (A, B) and CRY-negative s-CPDN3As (C, D). **(B’, D’)** Comparison of average Tb between experimental flies (E), *Gal4* control <u>(G),</u> and *UAS* control (U) during M bout in LD. Same statistical analysis as Fig. 2 was used. **(E, F)** Images of *spl-CRY-neg-LNd-Gal4* (*w;VT040042.AD;R31C03.DBD*) (E) and *spl-s-CPDN3A-Gal4* (*w;R14F03.AD;R31B03.DBD*) (F) crossed with *UAS-myrGFP* in green. Clock neurons were immunostained with anti-PER antibody (cyan). CRY-positive neurons were immunostained with anti-CRY antibody (magenta). The yellow dashed line indicates the midline of the brain.

Like the other CRY-negative oscillators, flies with TIM-depleted CRY-negative LNds lost both M and E peaks in males (Fig. 4A), while females lacked only the M peak (Fig. S5A) in DD. Both male and female flies, however, displayed lower body temperatures during the M bout and robust E rise (Fig. 4B, 4B′, S5B, and S5B′) in LD. Next, we focused on CRY-negative s-CPDN3As. In DD, both male and female flies with TIM-depleted CRY-negative s-CPDN3As lost the M peak but not E peak (Fig. 4C and S5C). In LD, males had a lower body temperature than the control group during the M bout in LD (Fig. 4D), although this difference was not statistically significant (Fig. 4D’). On the other hand, females exhibited statistically significant lower body temperatures during the M bout in LD (Fig. S5D and S5D’). These results suggest that CRY-negative LNds and s-CPDN3As are also involved in regulating morning BTR.

### CRY-negative oscillators control BTR in the morning: M bout in LD and M peak in DD

We demonstrated that TIM-depletion in each specific CRY-negative clock neurons led to a loss of a robust M bout in LD and M peak in DD, suggesting that CRY-negative clock neurons are responsible for BTR during the morning. Consequently, we investigated whether the clock in CRY-negative clock neurons is sufficient to restore BTR. First, we expressed PER in *per^01^* mutants using either the *Clk856-Gal4* or the *tim-Gal4* driver, which is the general Gal4 driver expressed in the most clock neurons. PER expression in all clock neurons (*Clk856-Gal4* and *tim-Gal4*) successfully restored a strong E rise in LD (Fig. 5A and B) and E peaks in DD (Fig. 5D, E). M peak in DD was restored with PER expression in all clocks using *Clk856-Gal4* (Fig. 5D). Notably, PER expression in all clock neurons was more effectively restored using *Clk856-Gal4* than *tim-Gal4*, likely due to differences in their expression levels across clock neurons. On the other hand, the *per^01^* flies that were made to express PER only in the CRY-negative neurons using the *tim-Gal4*;*Cry-Gal80* drivers exhibited a restored M bout in LD and M peak in DD, but neither E rise in LD nor E peak in DD (Fig. 5C and F). Thus, our data showed that CRY-negative neurons control morning BTR—M bout in LD and M peak in DD—which are different from the mechanisms controlling evening BTR—E rise in LD and E peak in DD (Fig. 6). Given that CRY-negative neurons contribute little to sleep–wake regulation, our results reveal their functional importance in BTR, highlighting that BTR and sleep–wake cycles are regulated through distinct mechanisms (Fig. S6).

**Figure 5:**
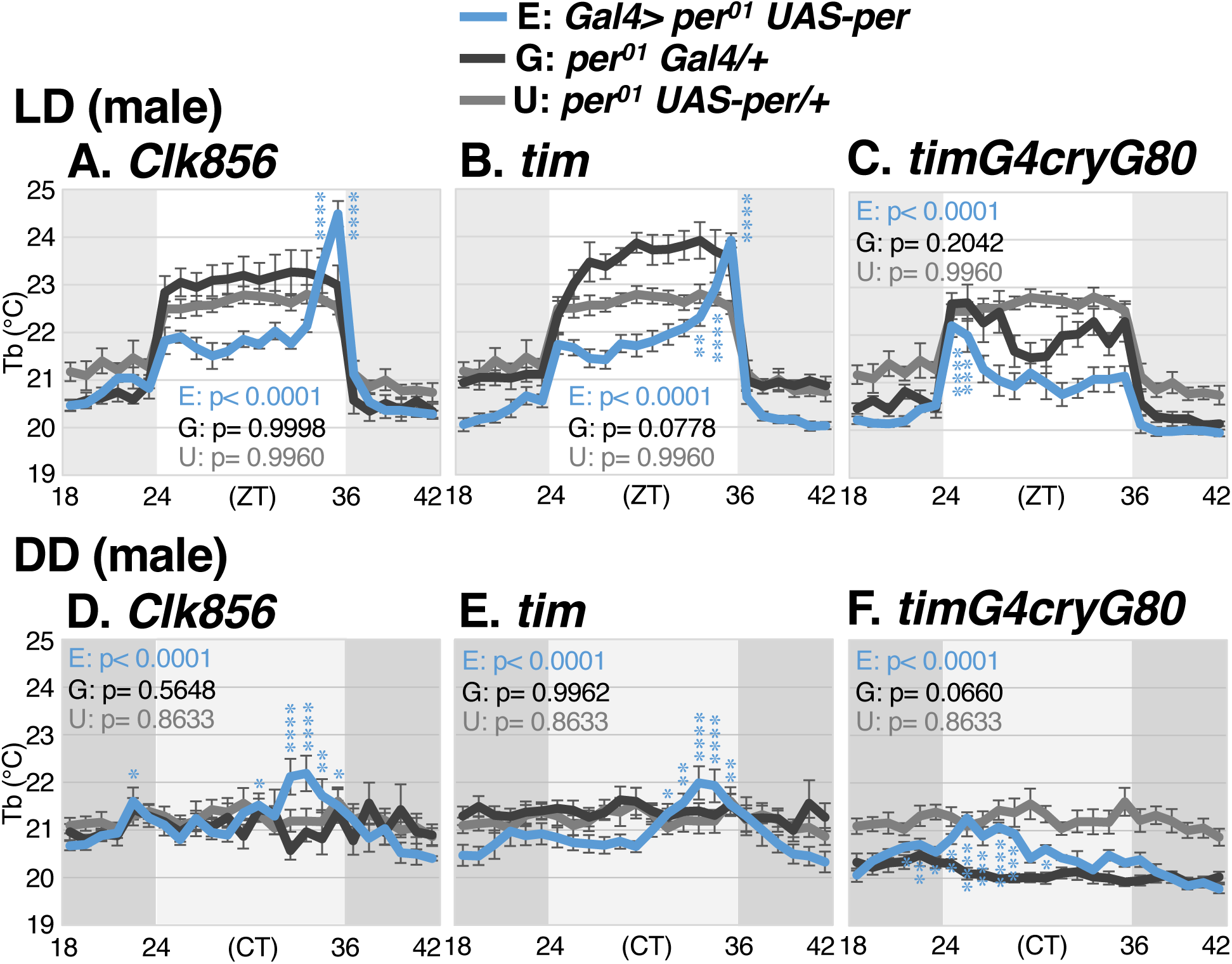
CRY-negative clock neurons restore M bout in LD and M peak in DD. **(A-F)** PER is selectively expressed in *per^01^* mutants to rescue BTR phenotype in all clock neurons (*Clk856-Gal4* (A, D), *tim*-*Gal4* (B, E)) and CRY-negative neurons (*Clk856-Gal4; Cry-Gal80*) (C, F) (each in blue). They were compared to *per^01^; Gal4* control (black) and *per^01^; UAS-per* control (gray). Rescue flies were tested in LD (A-C) and DD conditions (D-F). Same statistical analysis as Fig. 2 was used.

**Figure 6:**
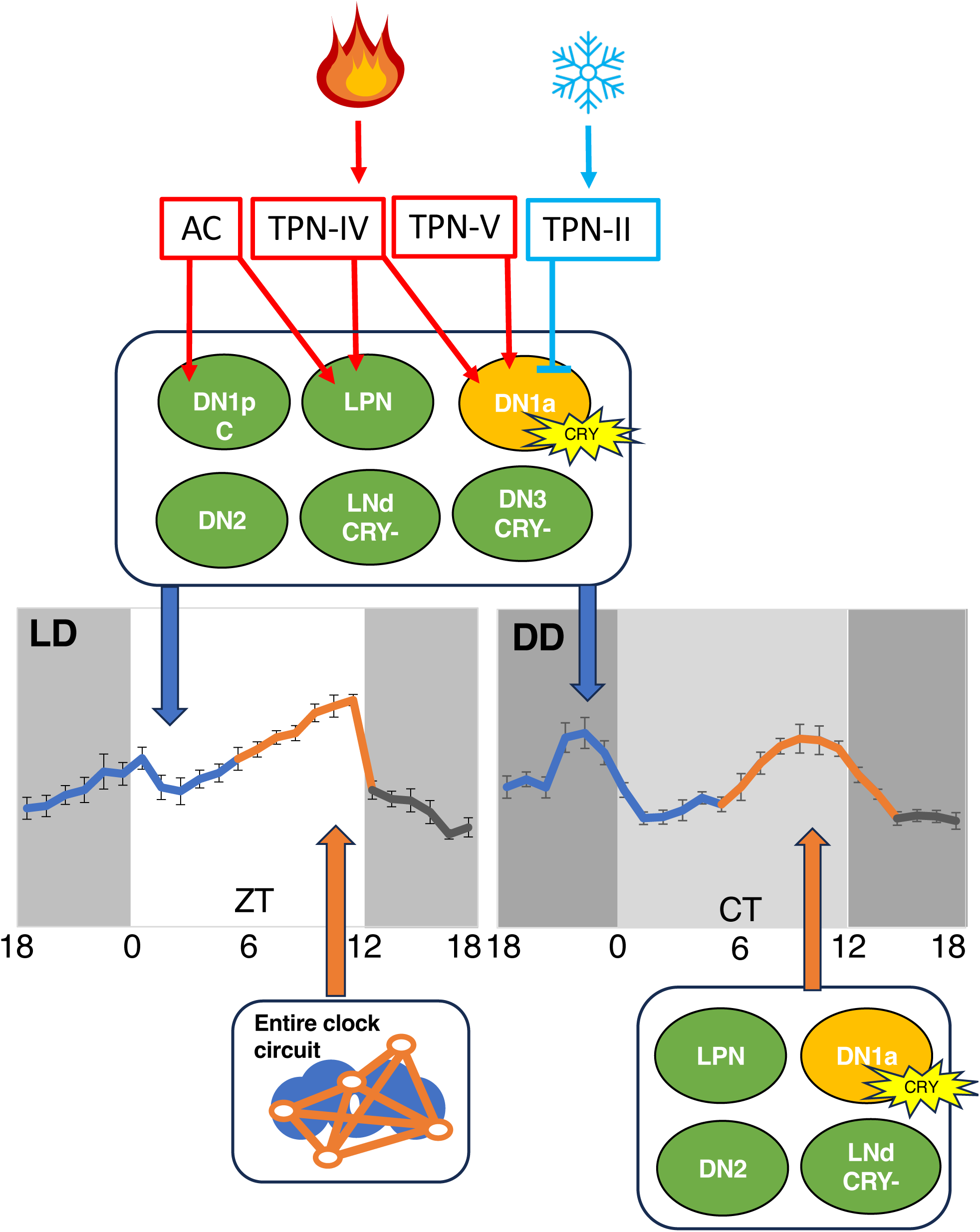
Schematic of the circuitry regulating BTR. DN1pCs, LPNs, and DN1as interact with the temperature processing neural circuits. DN1as also receive information from cold-sensing neurons. In addition to DN1pCs, LPNs, and DN1as, DN2s, CRY-negative LNds, and CRY-negative s-CPDN3As are all responsible for M bout and M peak (blue arrows in both LD and DD). E rise (orange arrow in LD) is regulated by all clock neurons and E peak (orange arrow in DD) is regulated by LPNs, DN1as, DN2s, and CRY-negative LNds. Green = CRY-negative clock neurons. Yellow = CRY-positive clock neurons.

## DISCUSSION

Body temperature rises to prepare animals for activities such as hunting and foraging. As the day progresses, it continues to increase and ultimately peaks in the evening. Many animals prefer warmer temperatures before sleep, while the subsequent decline in body temperature serves as an important cue for sleep initiation (*43–45*). Although the daily rhythm of body temperature (BTR) exhibits a conserved pattern observed across both endotherms and ectotherms, the mechanisms underlying the morning and evening BTR remain poorly understood. Here, we developed a machine learning-based assay, THERM-D, and found that CRY-negative clock neurons regulate morning BTR differently from evening BTR, suggesting that the body temperature increase in the morning and peak in the evening are controlled by different mechanisms. This finding also highlights that BTR is controlled separately from the sleep-wake cycles.

### THERM-D, a novel machine learning-based assay, enables continuous monitoring of BTR for over 24 hours

Locomotor activity behavior assays, which are commonly used to study circadian rhythms in *Drosophila,* have long served as a powerful, elegant, and efficient approach to uncover the fundamental importance of circadian rhythms. These studies have revealed many key regulatory mechanisms, several of which are evolutionarily conserved across animals (*46, 47*). Despite BTR being one of the strongest rhythms, body temperature and sleep–wake cycles are regulated independently, indicating that insights gained from locomotor activity behavior assays cannot fully explain the mechanisms underlying BTR. Given that BTR is an evolutionarily conserved and fundamental physiological mechanism that regulates animal homeostasis, establishing a dedicated model system for BTR was necessary.

We previously used TPR assays and discovered that *Drosophila* exhibit BTR (*18*). We found that the mechanisms controlling *Drosophila* BTR are different from the mechanisms controlling the sleep-wake cycle (*13*). Furthermore, we revealed that the G-protein coupled calcitonin receptor plays an important role for both *Drosophila* and mammalian BTR (*48*). However, we were concerned about the discontinuous nature of TPR assays that disturbed the flies’ sleep during night periods. Because body temperature is strongly associated with sleep, a new assay was needed to analyze BTR without disturbing sleep-wake cycles.

The major limitation of the TPR assays was that food could not be supplied to the apparatus, restricting fly survival to only a few hours. Starvation also induced a strong behavioral bias: hungry flies preferentially selected lower temperatures, likely as a strategy to conserve energy (*29*). As a result, reliable body temperature measurements could not be obtained beyond ∼1 hour. To construct a full 24-hour BTR curve, we had to perform TPR assays at eight separate time points using new sets of flies each time (*13*). At least five replicates were needed per time point, requiring a minimum of 40 independent experiments (Fig. S1A). Since each assay used 20–30 flies, a single 24-hour curve required a minimum of ∼800–1,200 flies per genotype. Because of this high demand, males and females were not evaluated separately.

Our development of the novel assay THERM-D set to address these limitations (Fig. 1A). By supplying food within the apparatus, flies remain satiated, eliminating starvation-induced cooling and enabling long-term, physiologically relevant measurements. With only 12–16 flies per chamber, their body temperature can be continuously tracked for 24 hours. A full dataset now requires only 72–96 flies (n = 6) or 108-144 flies (n=9), allowing us to separate sexes and directly compare behavioral differences between them.

By overcoming these constraints, THERM-D enables robust, continuous, and physiologically relevant monitoring of innate BTR phenotypes. Using THERM-D, we newly revealed that *Drosophila* exhibit morning and evening BTR throughout the day. In DD, they show robust bimodal morning and evening peaks (Fig. 1G). In LD, male flies show significant body temperature increase before lights on (Fig. 1B and C). Therefore, flies might be able to anticipate the morning and start to increase their body temperature in preparation for the coming dawn, which is the same as human BTR. We demonstrated that morning and evening BTR are separately regulated by the specific clock neurons (see below). These findings were able to be achieved by continuous and high-resolution data analysis using THERM-D.

### Clock neurons that have close interactions with temperature-sensing circuits regulate morning body temperature

Emerging results suggest a relationship between the thermoregulatory network and CRY-negative neurons. DN2s alter neuronal activity in response to rising or falling temperatures (*49*). LPNs receive inputs from ACs and TPN-IV and are involved in temperature-dependent sleep (*27*). DN1ps receive inputs from ACs. Among them, DN1pC neurons receive direct projections from ACs, suggesting that DN1pCs are the main targets of AC-derived input (*26, 28*). Here, using newly identified *Gal4* lines that drive a subset of CRY-negative neurons, we were able to reveal that several CRY-negative neurons determine BTR in the morning (Fig. 6). Moreover, we showed that DN1as are the critical core clock for BTR. DN1as are CRY-positive neurons and receive inputs from cold-processing neurons (TPN-II) and warm-processing neurons (TPN-IV and TPN-V) (*27*) (Fig. 6). Thus, the relationship between clock neurons and circadian temperature regulation appears to be crucial for driving the body temperature rise in the morning. This may explain why temperature-sensing circuits are directly linked to circadian circuits: the circuits work together to establish body temperature, fundamental to an animal’s temperature homeostasis.

### Distinct mechanisms regulate the morning and evening BTR

We showed that ablation of clocks in CRY-negative clock neurons caused flies to maintain a lower body temperature from late night through early morning under the LD condition, resulting in an abnormal M bout. However, they still showed a robust increase of their body temperature toward the E rise, suggesting that the mechanisms controlling the M bout and E rise of the BTR are distinct (Fig. 6).

We found different BTR patterns in LD and DD. In LD, flies show two different temperature increase periods, M bout and an E rise, whereas in DD they show dual, equivalent-looking M peak and E peaks. The key question is whether the M bout and E rise observed in LD correspond respectively to the M peak and E peak observed in DD. Flies with clock-disrupted CRY-negative neurons—LPNs, DN1pCs, DN2s, LNds (CRY-), and s-CPDN3As (CRY-)—along with CRY-positive DN1as all displayed abnormal M bout and M peak. These parallel phenotypes support the idea that M bout and M peak are governed by the same mechanisms. On the other hand, abnormal E rise was only observed when clocks of all clock neurons were ablated, while clock ablation in LPN, DN2, DN1a, or LNd (CRY-) eliminated E peak (Fig. 6). The data suggest that the mechanisms underlying E rise in LD and E peak in DD differ. Altogether, these claims support the idea that distinct mechanisms regulate the morning and evening BTR.

### DN2s and DN1as are core clock neurons for BTR

The clock ablation in either DN1as or DN2s showed no BTR in DD, suggesting that DN1as and DN2s are core clock neurons for BTR (Fig. 2C and E). However, clock ablation in either DN1as or DN2s still showed a complete E rise in LD (Fig. 2D and F). Even when the clocks in both DN1as and DN2s were knocked out simultaneously using *tim-Gal4, Pdf-Gal80*, or *CrzR-Gal4* (Fig. S3), the flies still exhibited E rise in LD. These results suggest that DN1as and DN2s are not the sole clock neurons driving the E rise; instead, multiple classes of clock neurons may be contributing. Notably, our previous TPR assay showed that TIM depletion in DN2s or DN1as eliminated daytime temperature increases (*33*). This discrepancy may reflect methodological differences between the TPR assay and THERM-D assay. One possibility is that in contrast to the light exposure for 30 minutes in the previous TPR assay apparatus, the flies in THERM-D apparatus are exposed to the light continuously for hours during the daytime in LD, which drives a robust temperature rise without the clock functions in DN2s and DN1as.

Furthermore, we demonstrated that clock depletion of either DN1a or DN2 results in a lower body temperature during M bout in LD (Fig. 2D’ and 2F’). However, knocking out TIM in both DN2s and DN1as simultaneously using *Clk856-Gal4* (Fig. 2B and 2B’), *CrzR-Gal4* (Fig. S3B and S3B’), or *tim-Gal4; Pdf-Gal80* (Fig. S3D and S3D’) caused no significant change in body temperature during M bout. These results suggest that the DN1a and DN2 circuits may interact reciprocally. For instance, if both DN1a and DN2 circuits are disrupted simultaneously, a change in temperature might not occur. If either DN1a or DN2 circuit is disrupted, the remaining circuit could abnormally control body temperature, resulting in a lower body temperature. Thus, the DN1a and DN2 circuits potentially influence one another, which could be important for regulating body temperature.

### BTR and sleep–wake cycles are controlled by different neuronal mechanisms

The independence of BTR regulation from sleep-wake cycles has been known in humans. Humans BTR fluctuates even under restricted locomotor activity (*50, 51*). Moreover, BTR and locomotor activity rhythms can be experimentally dissociated, a phenomenon known as spontaneous internal desynchronization (*52*). Previously, we demonstrated in flies that BTR and sleep-wake cycles are governed by distinct neural circuits (*17, 18*). Our THERM-D findings further support this concept by revealing that CRY-negative oscillators, which only play a minor role in regulating sleep-wake cycles, underlie the M bout in BTR (Fig. S6).

We wonder why CRY-negative oscillators are necessary for the M bout. Unlike CRY-negative neurons, CRY-positive clock neurons can directly respond to environmental light–dark cycles through the photoreceptive function of CRY, allowing their clocks to be rapidly entrained to external light cues (*53*). In contrast, CRY-negative neurons lack this direct light sensitivity and instead likely rely on indirect input from CRY-positive neurons or the visual system. Notably, CRY-negative neurons are more likely to be entrained by temperature cycles than by light–dark cycles (*25*). For this reason, the circuits regulating BTR may deliberately exclude direct light input. Such separation could explain the fundamental differences between the regulation of BTR and sleep–wake cycles.

Another mechanistic difference between BTR and sleep lies in their response to acute light exposure. In response to light, flies typically show simultaneous increases in both locomotor activity and body temperature (*31, 54*), suggesting that the acute rise in body temperature might simply result from increased locomotor activity. However, this is not always the case. For example, when TIM or PER is knocked out in LPNs, flies still exhibit robust locomotor activity rhythms with distinct M and E peaks (*55*). In contrast, TIM depletion in the LPNs abolishes the light-driven increase in body temperature during the M bout, resulting instead in a lowered body temperature during this period (ZT 18–24 through ZT 0–6). This indicates that the rise in body temperature following light exposure is not solely due to increased locomotor activity. Together, these findings further support the notion that BTR and sleep–wake cycles are governed by distinct regulatory mechanisms.

Our development of THERM-D has been pivotal in uncovering specific mechanisms underlying body temperature rhythms. BTR is distinguishable from other circadian behaviors and therefore holds much to be discovered. THERM-D will continue to shed light upon new mechanisms governing circadian temperature homeostasis.

**Figure S1: Previous TPR assay used to monitor *Drosophila* BTR and long-term BTR in DD using THERM-D**

**(A)** *Drosophila* temperature preference rhythm (TPR) assay. Flies are applied into a 6-lane plexiglass chamber atop a metal apparatus with Peltier devices maintain a temperature gradient of 18 to 32° C. Flies are placed in the apparatus for 30 minutes to document BTR at the given ZT timepoint. Experiments are repeated more than 5 times per ZT and manually analyzed. **(B)** Graph of continuous, 3-day *w^1118^* BTR in DD conditions using THERM-D. One-way ANOVA and post hoc Dunnett’s test were used for comparison to the lowest Tb during the whole experiment (CT 12-74) in DD. ∗∗∗∗p < 0.0001, ∗∗∗p < 0.001, ∗∗p < 0.01, or ∗p < 0.05 is shown. The bar under the graph represents the period we examined the BTR in this study: from CT 18 to 42.

**Figure S2. BTR of female flies with clock disruption in all clock neurons, DN2s, DN1as, or LNvs**

**(A-H)** Comparison of BTR between clock-disrupted flies (orange) and control flies (*Gal4* (black) and *UAS* (gray)) in LD and DD. Clocks were disrupted via targeted *tim*-CRISPR expression in all clock neurons (A, B), DN2s (C, D), DN1as (E, F), and LNvs (G, H). **(B’, D’, F’, H’)** Comparison of average Tb between experimental flies (E), *Gal4* control (G), and *UAS* control (U) during M bout in LD. Same statistical analysis as Fig. 2 was used. **(D’’)** Secondary analysis of BTR in LD of TIM-depleted DN2s, comparing average Tb during first half (ZT18-24) and second half (ZT0-6) of M bout between experimental flies (E), *Gal4* control (G), and *UAS* control (U). In D’’, ANOVA and post hoc Dunnett’s test were used for comparison to the lowest Tb during the first half (ZT 18-24) and second half (ZT 0-6) of M bout in LD.

**Figure S3. Dorsal clock neurons regulate BTR.**

**(A-D)** Comparison of clock-disrupted male flies using *tim*-CRISPR—DN1a and DN2s (A, B) and all dorsal clock neurons (C, D) in blue, *Gal4* control in black, and *UAS* control in gray, tested in DD and LD.

**(B’, D’)** Temperature comparison during M bout for LD between experimental (E), *Gal4* control (G), and *UAS* control (U). Same statistical analysis as Fig. 2 was used.

**(E)** Image of *CrzR-RB-Gal4* crossed with *UAS-myrGFP*. CrzR+ neurons are labeled with GFP (green). Clock neurons were immunostained with anti-TIMELESS (TIM) antibody (magenta).

(**F, G**) Temperature comparison of entire 24-hour DD result between experimental—DN2 (F) and LPN (G) in blue, *Gal4* control in black, and *UAS* control in gray. Comparison of average Tb between experimental (E), *Gal4* control (G), and *UAS* control (U) during entire 24-hour DD result (CT 18-24 and 0-18). ANOVA and post hoc Dunnett’s test were used for comparison between the experimental flies and Gal4 control or UAS control. ∗∗∗∗p < 0.0001, ∗∗∗p < 0.001, ∗∗p < 0.01, or ∗p < 0.05 is shown.

**Figure S4: BTR of female flies with clock disruption in CRY-negative LPN and DN1pC**

**(A-D)** Comparison of BTR between flies with targeted *tim* knockouts (orange) and control flies (*Gal4* (black) and *UAS* (gray)) in DD and LD. Clocks were disrupted via targeted *tim*-CRISPR expression in LPNs (A, B) and DN1pCs (C, D).

**(B’, D’)** Average temperature comparison of M bout in LD between experimental (E), *Gal4* control (G), and *UAS* control (U). Same statistical analysis as Fig. 2 was used.

**Figure S5: BTR of female flies with clock disruption in CRY-negative LNd and s-CPDN3A**

**(A-D**) Comparison of BTR in DD and LD between flies with clock disruption (orange) and control flies (*Gal4* (black) and *UAS* (gray)). Clocks were disrupted via *tim*-CRISPR expression in CRY-negative LNds (A, B) and CRY-negative s-CPDN3As (C, D).

**(B’, D’)** Comparison of average Tb between experimental (E), *Gal4* control (G), and *UAS* control (U) during M bout. Same statistical analysis as Fig. 2 was used.

**Figure S6: A comparison of neural circuitry underlying locomotor activity rhythms and BTR in LD.**

**(A)** LNvs and DN1ps regulate M peak; 5^th^-LNv, LNds, and DN1ps regulate E peak in locomotor activity rhythms. **(B)** DN1a, DN2, LPN, DN1pC, CRY-negative LND, and CRY-negative s-CPDN3As regulate M bout, and the entire clock circuit regulates E rise in LD.

## Material and Method

### Fly Strains

*w^1118^* (BDSC:5905; RRID:BDSC_5905), *Spl-DN1a-Gal4* (*23E05-p65.AD* (BDSC_70601) and *R92H07-Gal4.DBD* (BDSC_70004)), *Pdf-gal4* (BDSC_6900), *Spl-LPN-Gal4* (*w; R11B03-p65.AD; R65D05-Gal4.DBD* (BDSC_89189)), *Spl-DN1pC-Gal4* (*R51H04.AD; VTO26178.DBD* (BDSC_89174)), *Spl-CRYnegativeLNd-Gal4* (*VT040042AD;R31C03.DBD* (BL88484)), *Spl-s-CPDN3A-Gal4* (*R14F03.AD;R31B01.DBD* (BDSC_89061)), *CrzR-RB-Gal4* (BDSC_84621), *UAS-GFP S65T* (BDSC_1522) were provided by the Bloomington Drosophila Stock Center. *wCS, per^01^, tim^01^, UAS-sgRNA-tim^3x^; UAS-Cas9.2, Clk856-Gal4* were kindly shared from Dr. William Ja, Dr. Paul H Taghert, Dr. Patrick Emery, Dr. Mimi Shirasu-Hiza, and Dr. Orie Shafer, respectively.

### Other flies

*Clk9M-Gal4; PdfGAL80* (*13*)*, Tim-Gal4; Pdf-GAL80* (*37*)*, Tim(UAS)-Gal4* (*56*) *Cry-GAL80* (*37*), UAS-myrGFP (*57*)

### THERM-D Assay

All flies were raised in the same 12-hour light-dark (LD) incubator at 25°C and 60% humidity with a fluorescent light intensity of 600-1000 lux. Male and female flies were collected in the same vials at age day 0-4 and were separated 1-2 days later, on day 1-5. This ensured all females were mated and allowed for sexual dimorphic behaviors to become apparent.

The apparatus consists of a metal platform and 6-lane plexiglass cover. A temperature gradient, ranging from 18-32°C, is generated on the apparatus and recorded through probes. Food (5% sucrose and 2% agar in deionized water) is placed on the cold end of the apparatus, which is kept below 19°C, to ensure the flies can survive for the duration of the experiment. The flies normally avoid below 19 °C because they find it unpleasant. Therefore, the flies can only remain around the food for a very short time. Each genotype was tested in both 12-hour LD cycles and constant darkness (DD) conditions in an environmental room maintained at 25°C and 35-50% humidity. Two to four hours prior to the light off (ZT 12), flies were placed into the chambers on the apparatus. In this setup, approximately 15 flies per lane were freely allowed to select their body temperature (Tb) over the course of 2-3 days. Males and females were tested separately. The photos and temperature information were taken every hour to record the Tb using a timer-controlled Canon EOS Rebel T100 DSLR Camera with flash on and a 4-probe temperature tracker.

THERM-D assays were performed continuously for 2-3 days with the same flies. The photos taken from the middle of the first night (ZT/CT 18-24), first day (ZT/CT 0-12), and second night (ZT/CT 12-18) were analyzed using a Python-based machine-learning algorithm (please see the details in another file for Supplemental information). Because the Tb sometimes varies among different fly lines, the Tb of *Gal4/UAS* flies was always compared with that of *Gal4/+* or *UAS/+* control flies.

#### IMMUNOHISTOCHEMISTRY

Male flies were entrained in LD12:12 at 25 °C for at least 3 days. Whole flies were sampled shortly before lights on (ZT0) to allow CRY to accumulate overnight. The flies were fixed in 4% paraformaldehyde in phosphate-buffered saline (PBS) with 0.1% Triton X-100 (PBS-T) for 3h at room temperature. Fixed flies were washed three times with PBS before dissections. The brains were blocked overnight at 4°C in PBS-T containing 5% normal donkey serum and subsequently incubated in primary antibody solutions at 4 °C for 48 h. The following primary antibodies were used (Chicken anti-GFP 1:1000 (Abcam, RRID: AB_300798), Goat anti-PER 1:200 (Santa Cruz Biotechnology, RRID: AB_654018), Rabbit anti-dCRY 1:1000 (Todo, T RRID: AB_2314242)). Following six washes with PBS-T, the brains were incubated in highly cross adsorbed secondary antibodies (all diluted 1:200; Thermo Fisher Scientific; Alexa Fluor 488 donkey anti-chicken, Alexa Fluor 555 donkey anti-rabbit, Alexa Fluor 647 donkey anti-goat) at 4°C overnight. Lastly, the samples were washed six times in PBS-T and mounted in Vectashield mounting medium (Vector Laboratories, Burlingame, CA, USA).

Samples were scanned with a Leica TCS SP8 confocal microscope equipped with a hybrid detector. A white light laser (Leica Microsystems, Wetzlar, Germany) was used for excitation. A 20-fold glycerol immersion objective (0.73 NA, HC PL APO, Leica Microsystems, Wetzlar, Germany) was used for wholemount scans, resulting in confocal stacks with 2048 × 1024 pixels with a maximal voxel size of 0.3 × 0.3 × 2 µm and an optical section thickness of 3.12 µm. For noise reduction, a frame average of 3 was used. Images were acquired using Leica Application Suite X (v3.5.7.23225) and analyzed using Fiji.

##### CrzR-Gal4

Immunostainings were performed as described previously (*58*). The following antibodies and dilutions were used: Rat anti-TIM (1:3000; kindly provided by Jadwiga Giebultowicz) (*24*), Chicken anti-GFP (1:1000; Rockland, Cat# 600-901-215), Goat anti-chicken-Alexa 488 nm (1:1000; Life Technologies, Cat# A11039) and Goat anti-rat-Cy3 (1:1000; Millipore, Cat# AP136C) antibodies. Images were taken using laser scanning confocal microscopes (Olympus FV3000, Olympus, Tokyo, Japan).

### Identification of split-Gal4 lines

Clock neuron subtypes were previously identified in the FAFB connectome (*26, 59*). Single reconstructions of CRY-negative neurons were queried in Codex (*60*) against the “FlyLight Annotator Gen1 MCFO” library (gen1mcfo.janelia.org) using brainCircuits.io (braincircuits.io). The obtained Color-MIP-Maps were screened for morphologically similar neurons. Candidate Gal4 lines were further queried for split-Gal4 parts in the FlyLight Raw collection for split-Gal4 combinations (flylight-raw.janelia.org). The expression patterns of split-Gal4 combinations were screened for morphologically similar neurons to clock neuron subtypes. Potential candidates were ordered at the Bloomington Drosophila Stock Center and crossed to a membrane-bound GFP reporter for anatomical validation. The brains were stained against PER and CRY for further validation. The identified lines are part of the split-Gal4 collections published in (*61*) and Dione et al. (in preparation).

### Quantification and statistical analysis

For the statistical analysis in BTR, Tb from ZT 0-12 in LD or CT 18-42 in DD were compared with one-way ANOVA, then Dunnett’s post hoc test was performed against the lowest Tb among ZT 0-12 in LD or CT18-42 in DD. For the statistical analysis in M bout, the average Tb from ZT 18-30 in *Gal4*/*UAS* flies was compared to that of *Gal4*/+ or *UAS*/+ control flies with one-way ANOVA and Dunnett’s post hoc test. All the statistical analysis were performed using GraphPad Prism10. All significance values are denoted in each graph.

## Supporting information

Suppl merged all

## ACKNOWLEDGMENTS

We are grateful to Drs. Mimi Shirasu-Hiza and the Bloomington *Drosophila* fly stock center for providing the fly lines; the Hamada lab members especially Alexis M Atherley, Jeff Mcgee, Gurbani Saini, Phoebe A Nicholas, Rasneet Grewal, Thy Pham, Benjamin E Falkenstein, Anthony Yap, for helping with experiments. This research was supported by JSPS KAKENHI grant 24K09534 to T.Y., NIH R01 grant GM107582, R21 grant

NS112890, R35 grant GM152154, R34 grant NS132843 to F.N.H, Eugene Cota-Robles Fellowship to O.M.L., and partly by DFG grant FO207/16-1.

## AUTHOR CONTRIBUTIONS

Conceptualization, F.N.H., T.G., and O.M.L.; Methodology, F.N.H., T.G., R.R., N.U., C.T.B., and J.L.; Investigation, T.G., O.M.L., N.R., A.F., K.R., M.G.A., R.C., V.Z.M., R.N., N.R., and C.F.; Writing – Original Draft, F.N.H., T.G., and O.M.L.; Writing –Review & Editing, F.N.H., T.G., O.M.L, N.R, C.H, and T.Y. Funding Acquisition, F.N.H., and T.Y.; Visualization, F.N.H., T.G., and O.M.L.; Resources, Y.U.; Supervision, F.N.H.

## DECLARATION OF INTERSTS

The authors declare no competing interests

## REFERENCES

1. J. Aschoff, Circadian control of body temperature. J. therm.Biol 8, 143–147 (1983).

2. K. Krauchi, How is the circadian rhythm of core body temperature regulated? Clin Auton Res 12, 147–149 (2002).

3. D. Weinert, Circadian temperature variation and ageing. Ageing Res Rev 9, 51–60 (2010).

4. R. Refinetti, M. Menaker, The circadian rhythm of body temperature. Physiol Behav 51, 613–637 (1992).

5. L. C. Lack, M. Gradisar, E. J. Van Someren, H. R. Wright, K. Lushington, The relationship between insomnia and body temperatures. Sleep Med Rev 12, 307–317 (2008).

6. J. F. Duffy, D. J. Dijk, E. B. Klerman, C. A. Czeisler, Later endogenous circadian temperature nadir relative to an earlier wake time in older people. The American journal of physiology 275, R1478–1487 (1998).

7. E. D. Buhr, S. H. Yoo, J. S. Takahashi, Temperature as a universal resetting cue for mammalian circadian oscillators. Science 330, 379–385 (2010).

8. J. Morf, U. Schibler, Body temperature cycles: gatekeepers of circadian clocks. Cell Cycle 12, 539–540 (2013).

9. K. Krauchi, The human sleep-wake cycle reconsidered from a thermoregulatory point of view. Physiol Behav 90, 236–245 (2007).

10. K. Krauchi, The thermophysiological cascade leading to sleep initiation in relation to phase of entrainment. Sleep Med Rev 11, 439–451 (2007).

11. S. S. Gilbert, C. J. van den Heuvel, S. A. Ferguson, D. Dawson, Thermoregulation as a sleep signalling system. Sleep Med Rev 8, 81–93 (2004).

12. F. N. Hamada et al., An internal thermal sensor controlling temperature preference in Drosophila. Nature 454, 217–220 (2008).

13. H. Kaneko et al., Circadian Rhythm of Temperature Preference and Its Neural Control in Drosophila. Current biology: CB 22, 1851–1857 (2012).

14. D. Giraldo, A. Adden, I. Kuhlemann, H. Gras, B. R. H. Geurten, Correcting locomotion dependent observation biases in thermal preference of Drosophila. Sci Rep 9, 3974 (2019).

15. R. D. Stevenson, Body size and limits to the daily range of body temperature in terrestrial ectotherms. Am. Nat., 102–117 (1985).

16. R. D. Stevenson, The relative importance of behavioral and physiological adjustments controlling body temperature in terrestrial ectotherms. The American Naturalist 126, (1985).

17. T. Goda, F. N. Hamada, Drosophila Temperature Preference Rhythms: An Innovative Model to Understand Body Temperature Rhythms. Int J Mol Sci 20, (2019).

18. T. Goda, Y. Umezaki, F. N. Hamada, Molecular and Neural Mechanisms of Temperature Preference Rhythm in Drosophila melanogaster. Journal of biological rhythms 38, 326–340 (2023).

19. J. Barrett, L. Lack, M. Morris, The sleep-evoked decrease of body temperature. Sleep 16, 93–99 (1993).

20. P. J. Murphy, S. S. Campbell, Nighttime drop in body temperature: a physiological trigger for sleep onset? Sleep 20, 505–511 (1997).

21. K. J. Fogle, K. G. Parson, N. A. Dahm, T. C. Holmes, CRYPTOCHROME is a blue-light sensor that regulates neuronal firing rate. Science 331, 1409–1413 (2011).

22. P. Emery, W. V. So, M. Kaneko, J. C. Hall, M. Rosbash, CRY, a Drosophila clock and light-regulated cryptochrome, is a major contributor to circadian rhythm resetting and photosensitivity. Cell 95, 669–679 (1998).

23. J. Benito, J. H. Houl, G. W. Roman, P. E. Hardin, The blue-light photoreceptor CRYPTOCHROME is expressed in a subset of circadian oscillator neurons in the Drosophila CNS. Journal of biological rhythms 23, 296–307 (2008).

24. T. Yoshii, T. Todo, C. Wulbeck, R. Stanewsky, C. Helfrich-Forster, Cryptochrome is present in the compound eyes and a subset of Drosophila’s clock neurons. The Journal of comparative neurology 508, 952–966 (2008).

25. T. Yoshii, C. Hermann, C. Helfrich-Forster, Cryptochrome-positive and - negative clock neurons in Drosophila entrain differentially to light and temperature. Journal of biological rhythms 25, 387–398 (2010).

26. N. Reinhard et al., Synaptic connectome of the Drosophila circadian clock. Nature communications 15, 10392 (2024).

27. M. H. Alpert, H. Gil, A. Para, M. Gallio, A thermometer circuit for hot temperature adjusts Drosophila behavior to persistent heat. Current biology: CB 32, 4079–4087 e4074 (2022).

28. X. Jin et al., A subset of DN1p neurons integrates thermosensory inputs to promote wakefulness via CNMa signaling. Current biology: CB 31, 2075–2087 e2076 (2021).

29. Y. Umezaki et al., Feeding-State-Dependent Modulation of Temperature Preference Requires Insulin Signaling in Drosophila Warm-Sensing Neurons. Current biology: CB 28, 779–787.e773 (2018).

30. T. L. Page, Masking in invertebrates. Chronobiology international 6, 3–11 (1989).

31. L. M. Head et al., The influence of light on temperature preference in Drosophila. Current biology: CB 25, 1063–1068 (2015).

32. R. Dubruille, P. Emery, A plastic clock: how circadian rhythms respond to environmental cues in Drosophila. Mol Neurobiol 38, 129–145 (2008).

33. S. C. Chen et al., Dorsal clock networks drive temperature preference rhythms in Drosophila. Cell Rep 39, 110668 (2022).

34. R. Delventhal et al., Dissection of central clock function in Drosophila through cell-specific CRISPR-mediated clock gene disruption. Elife 8, (2019).

35. M. Schlichting, M. M. Diaz, J. Xin, M. Rosbash, Neuron-specific knockouts indicate the importance of network communication to Drosophila rhythmicity. Elife 8, (2019).

36. F. Port, S. L. Bullock, Augmenting CRISPR applications in Drosophila with tRNA-flanked sgRNAs. Nat Methods 13, 852–854 (2016).

37. D. Stoleru, Y. Peng, J. Agosto, M. Rosbash, Coupled oscillators control morning and evening locomotor behaviour of Drosophila. Nature 431, 862–868 (2004).

38. K. M. Parisky et al., PDF cells are a GABA-responsive wake-promoting component of the Drosophila sleep circuit. Neuron 60, 672–682 (2008).

39. B. Grima, E. Chelot, R. Xia, F. Rouyer, Morning and evening peaks of activity rely on different clock neurons of the Drosophila brain. Nature 431, 869–873 (2004).

40. V. Sheeba et al., Large ventral lateral neurons modulate arousal and sleep in Drosophila. Current biology: CB 18, 1537–1545 (2008).

41. X. Tang et al., The role of PDF neurons in setting preferred temperature before dawn in Drosophila. Elife 6, (2017).

42. J. D. Ni et al., Differential regulation of the Drosophila sleep homeostat by circadian and arousal inputs. Elife 8, (2019).

43. E. C. Harding et al., A Neuronal Hub Binding Sleep Initiation and Body Cooling in Response to a Warm External Stimulus. Current biology: CB 28, 2263–2273 e2264 (2018).

44. E. C. Harding, N. P. Franks, W. Wisden, Sleep and thermoregulation. Curr Opin Physiol 15, 7–13 (2020).

45. E. C. Harding, N. P. Franks, W. Wisden, The Temperature Dependence of Sleep. Front Neurosci 13, 336 (2019).

46. O. T. Shafer, A. C. Keene, The Regulation of Drosophila Sleep. Current biology: CB 31, R38–R49 (2021).

47. O. T. Shafer, 25 years of Drosophila "Sleep genes". Fly 19, 2502180 (2025).

48. T. Goda et al., Calcitonin receptors are ancient modulators for rhythms of preferential temperature in insects and body temperature in mammals. Genes Dev 32, 140–155 (2018).

49. S. Yadlapalli et al., Circadian clock neurons constantly monitor environmental temperature to set sleep timing. Nature, (2018).

50. R. E. Smith, Circadian variations in human thermoregulatory responses. J Appl Physiol 26, 554–560 (1969).

51. P. H. Gander, L. J. Connell, R. C. Graeber, Masking of the circadian rhythms of heart rate and core temperature by the rest-activity cycle in man. Journal of biological rhythms 1, 119–135 (1986).

52. P. Lavie, Sleep-wake as a biological rhythm. Annu Rev Psychol 52, 277–303 (2001).

53. P. Emery et al., Drosophila CRY is a deep brain circadian photoreceptor. Neuron 26, 493–504 (2000).

54. B. Lu, W. Liu, F. Guo, A. Guo, Circadian modulation of light-induced locomotion responses in Drosophila melanogaster. Genes Brain Behav 7, 730–739 (2008).

55. C. Y. P. Guerrero, M. R. Cusick, A. J. Samaras, N. S. Shamon, D. J. Cavanaugh, The cell-intrinsic circadian clock is dispensable for lateral posterior clock neuron regulation of Drosophila rest-activity rhythms. Neurobiol Sleep Circadian Rhythms 18, 100124 (2025).

56. J. Blau, M. W. Young, Cycling vrille expression is required for a functional Drosophila clock. Cell 99, 661–671 (1999).

57. B. D. Pfeiffer et al., Refinement of tools for targeted gene expression in Drosophila. Genetics 186, 735–755 (2010).

58. A. Fukuda, A. Saito, T. Yoshii, Neurotransmitter and Receptor Mapping in Drosophila Circadian Clock Neurons via T2A-GAL4 Screening. Journal of biological rhythms 40, 491–497 (2025).

59. S. Dorkenwald et al., Neuronal wiring diagram of an adult brain. Nature 634, 124–138 (2024).

60. A. Matsliah, et al., Codex: Connectome Data Explorer. (2023).

61. G. W. Meissner et al., A split-GAL4 driver line resource for Drosophila neuron types. Elife 13, (2025).

