## Supplementary material for "THERM-D Uncovers Distinct Neural Mechanisms Separating Morning and Evening Body Temperature Rhythms in *Drosophila*": Suppl merged all

### Fig. S1

#### A. TPR assay

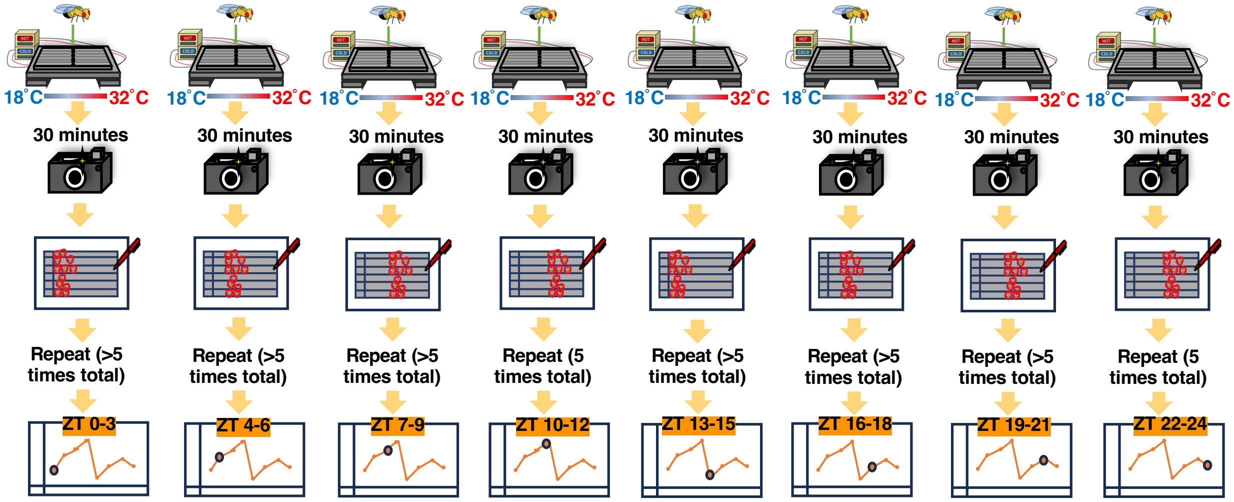


#### B. Continuous BTR in DD

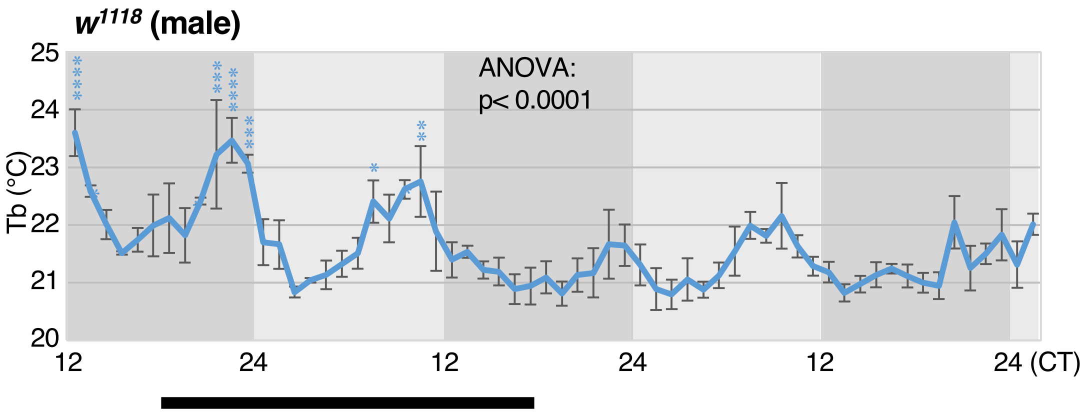

**Fig. S2**
**Female**

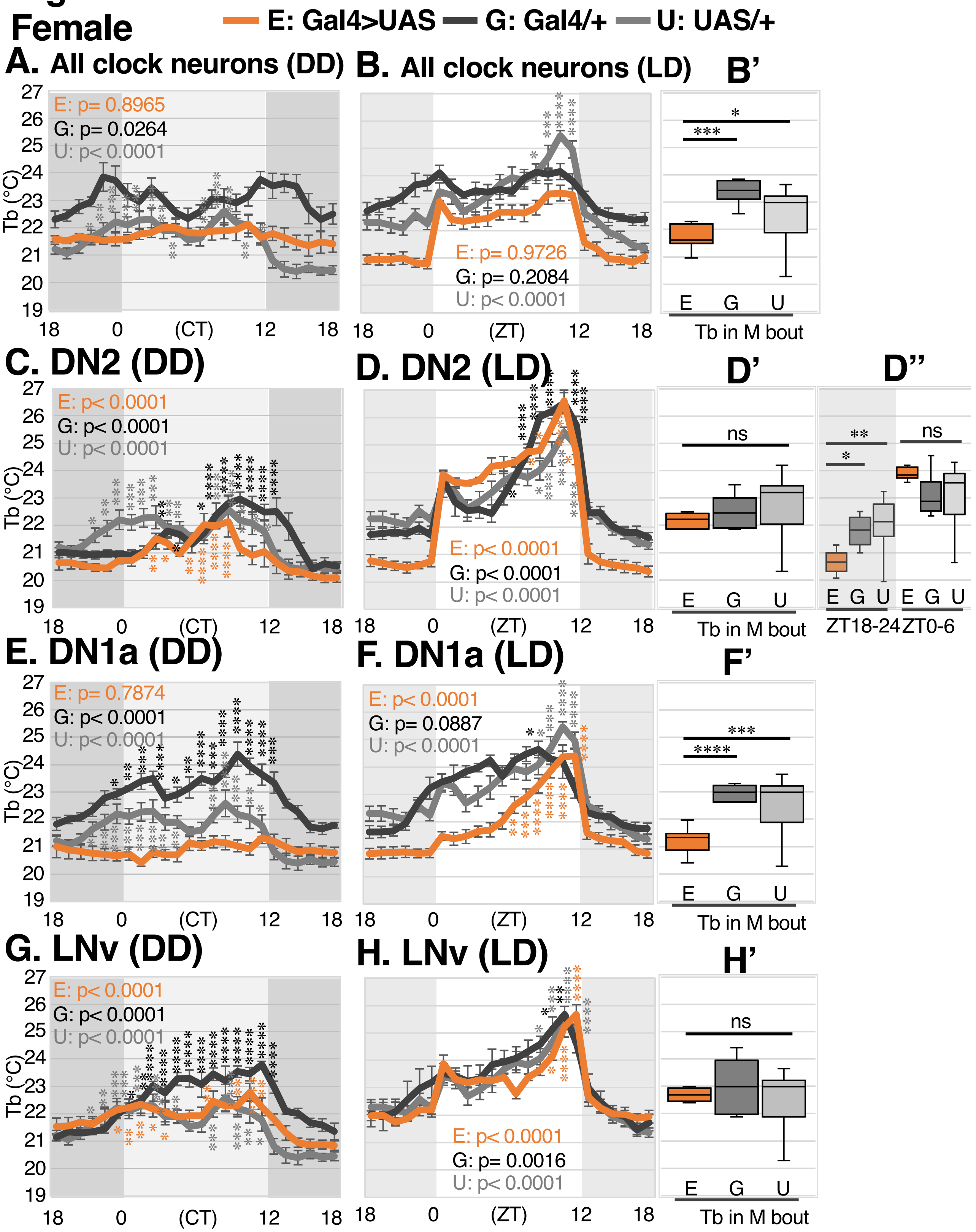

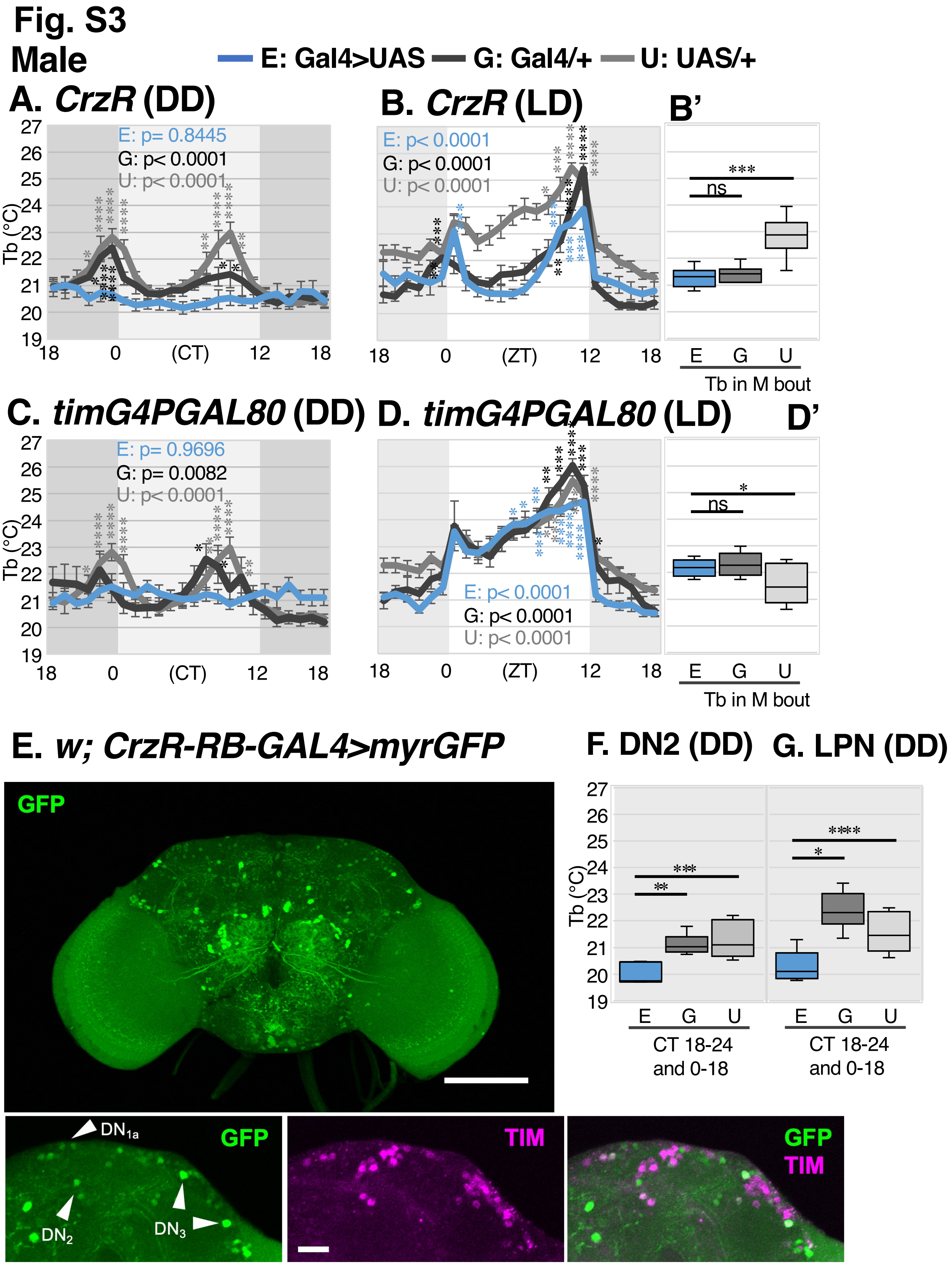
Fig. S3
Male
E: Gal4>UAS
G: Gal4/+
U: UAS/+
A. CrzR (DD)
E: p= 0.8445
G: p< 0.0001
U: p< 0.0001
Tb (°C)
(CT)
B. CrzR (LD)
E: p< 0.0001
G: p< 0.0001
U: p< 0.0001
(ZT)
B'
***
ns
E G U
Tb in M bout
C. timG4PGAL80 (DD)
E: p= 0.9696
G: p= 0.0082
U: p< 0.0001
D. timG4PGAL80 (LD)
E: p< 0.0001
G: p< 0.0001
U: p< 0.0001
D'
*
ns
Tb in M bout
E. w; CrzR-RB-GAL4>myrGFP
GFP
F. DN2 (DD)
G. LPN (DD)
CT 18-24 and 0-18
DN1a
DN2
DN3
TIM
GFP TIM

**Fig. S4**

**Female**

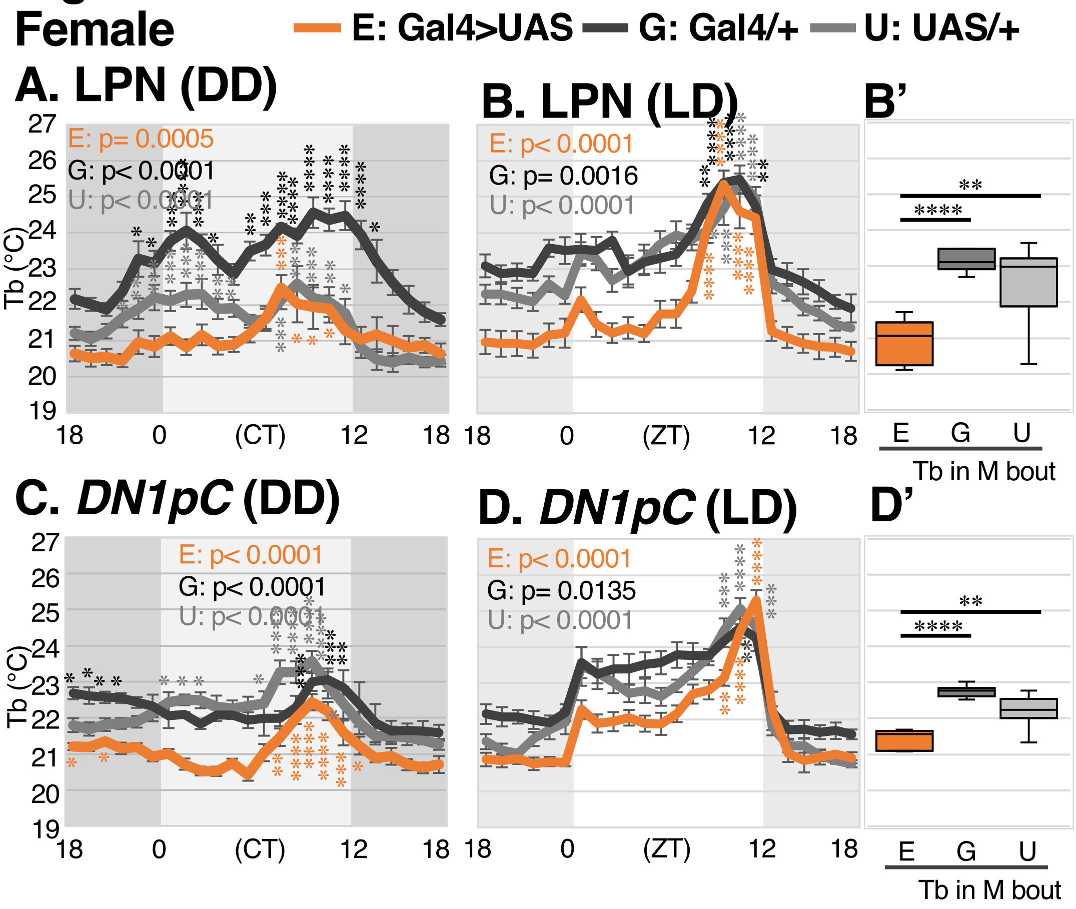

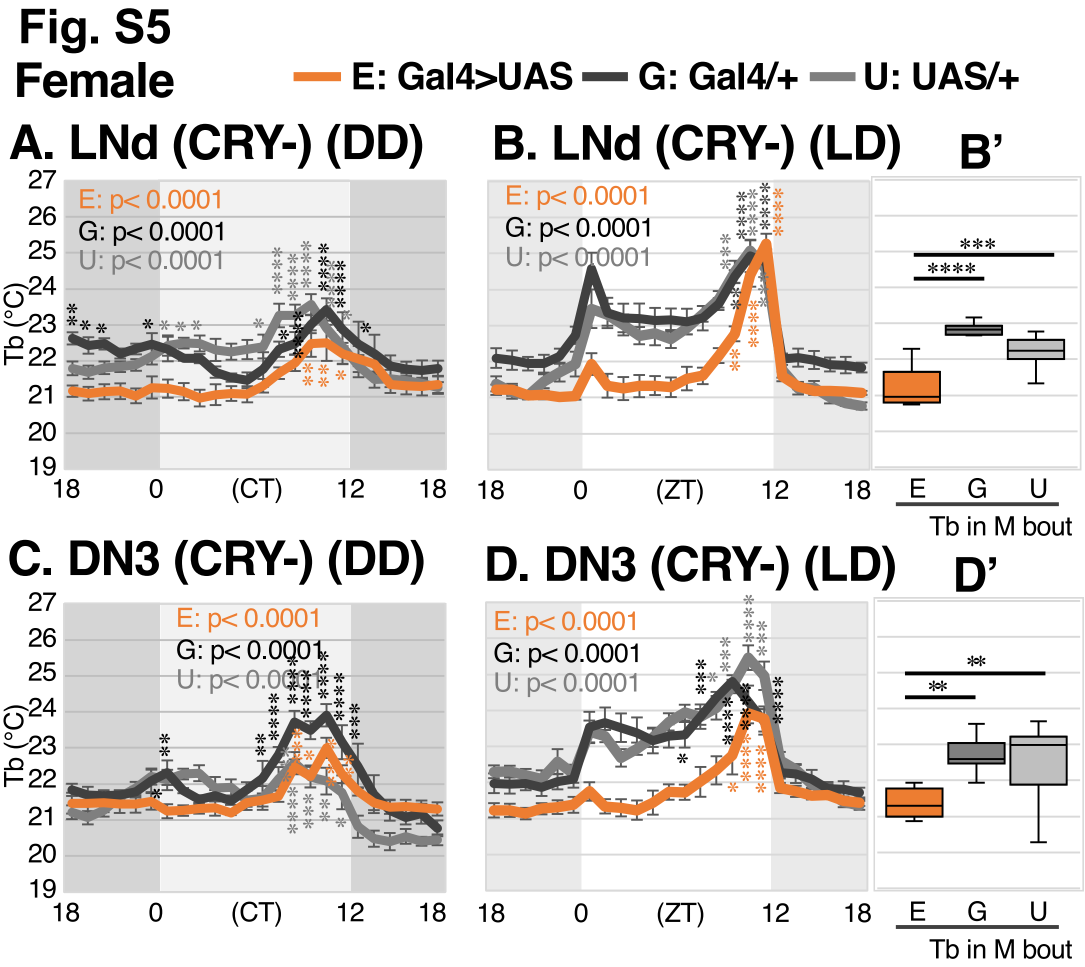
Fig. S5
Female
E: Gal4>UAS
G: Gal4/+
U: UAS/+
A. LNd (CRY-) (DD)
B. LNd (CRY-) (LD)
B'
E: p< 0.0001
G: p< 0.0001
U: p< 0.0001
Tb (°C)
(CT)
(ZT)
E G U
Tb in M bout
C. DN3 (CRY-) (DD)
D. DN3 (CRY-) (LD)
D'

#### Fig. S6

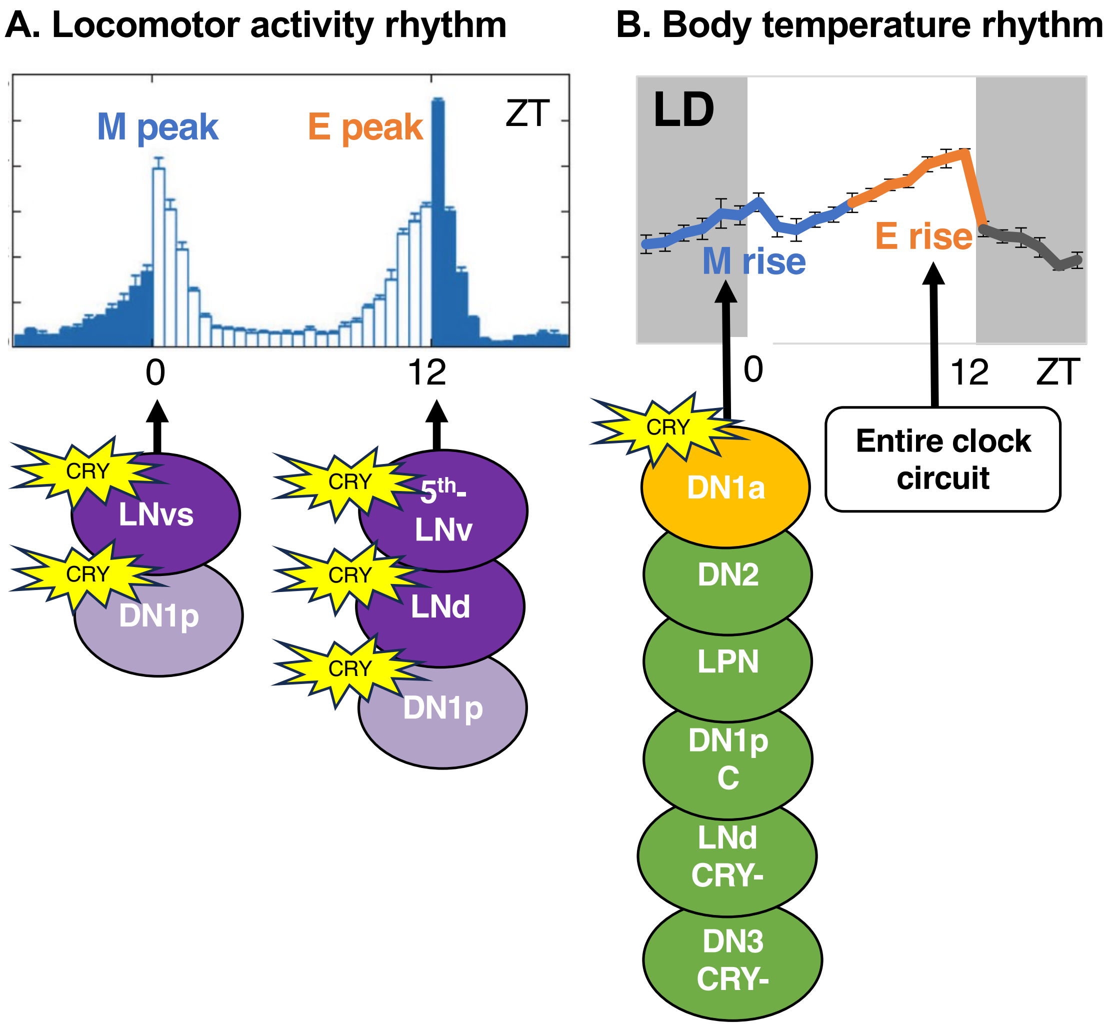
A. Locomotor activity rhythm
M peak
E peak
ZT
0
12
CRY
LNvs
CRY
DN1p
CRY
5th-LNv
CRY
LNd
CRY
DN1p
B. Body temperature rhythm
LD
M rise
E rise
0
12
ZT
CRY
DN1a
DN2
LPN
DN1p C
LNd CRY-
DN3 CRY-
Entire clock circuit

### Supplementary Material

This document contains additional information for the manuscript entitled "THERM-D Uncovers Distinct Neural Mechanisms Separating Morning and Evening Body Temperature Rhythms in *Drosophila"*

#### Image Processing Workflow

To expedite the analysis of our fly behavioral data on the temperature gradients, we developed a computational image processing workflow with the Python programming language. This workflow automates the process of locating and estimating temperatures for flies in photographic images of the experimental apparatus (such as in Fig. 1A). The workflow consists of three phases: 1) a registration phase, which rotates and crops each image to the area that contains flies; 2) a fly detection phase, which uses a machine learning model on the standardized image to estimate the location of each fly as (x, y) pixel coordinates, and converts these to physical coordinates (using the known physical size of the arena) and uses linear interpolation to estimate the temperature at each location; 3) a cleaning phase, which removes false positives and compiles lane-specific data. We designed the software to be robust against multiple varying image conditions including lighting, camera orientation, and background details that vary within and across experiments. Moreover, it allowed for data extraction from multiple experimental apparatuses that differ slightly in appearance.

##### Phase 1: Registration of Fly Arenas

The registration phase of the software workflow prepares the images for the machine learning model by correcting the orientation, removing any skew, standardizing the brightness, and cropping out irrelevant background details, so that the entire image consists of the fly arena. This phase has two parts, which we describe in the following paragraphs.

In Part 1, independently for each image, we estimate the locations of four registration marks (one orange, three green; see Fig. 1A) that we affixed to the apparatus at the four corners of the arena.

An image can be represented digitally by a rectangular array of small solid-colored squares called pixels (short for "picture elements"). By assuming a "color model," a model that describes how to represent colors with numerical coordinates, an image can thus be represented by an array of numbers. For example, in the popular "Red, Green, Blue" (RGB) color model, each color is represented by three numbers, called "channels," that indicate intensity of red, green, and blue light, respectively. With the RGB color model, an image can be represented by a 3-dimensional array of numbers. In our code for registering fly arenas, we use the "Hue, Saturation, and Value" (HSV) color model (Joblove & Greenberg 1978). The hue channel controls the perceived hue (from red to violet), the saturation channel controls the perceived colorfulness (from grayscale to full color), and the value channel controls the perceived brightness (from dark to light). The HSV color model is useful in image processing applications because it decouples hue, saturation, and brightness; in contrast, in the RGB color model, changing hue, saturation, or brightness typically requires changing all three channels.

The lighting conditions for each image vary, so we standardize the brightness of the image by applying adaptive gamma correction (AGC) (Rahman et al., 2016). AGC makes relatively dark parts of the image brighter and relatively bright parts of the image darker, which is beneficial for locating the registration marks. We chose AGC from several contrast enhancement methods because, for a sample of our images, it was best at standardizing brightness while preserving visual detail. To apply AGC, our code rescales the brightness value of each pixel based on the overall brightness of the image. Hue and saturation are left unchanged.

After AGC, our code searches for pixels in the image that match the expected HSV coordinates of the registration marks. Although AGC mitigates extreme variation, there is still some natural variation between images that can affect registration mark detection. To account for this, our code then searches for pixels within the range of HSV coordinates expected to correspond to registration marks. For example, for green registration marks, we search for all pixels with hues between 70 and 85 (each channel ranges from 0 to 255), saturation greater than 63, and brightness greater than 95. We set the lower bound on brightness for the orange registration marks adaptively at the 45th percentile of the image's brightness, since we found that a fixed lower bound for this case did not work well in practice. The result of this step is a matrix for each type of registration mark (green, orange) that respectively labels each pixel as likely (1) or unlikely (0) to be part of the respective registration mark (see Fig. 1B-C). These matrices are referred to as green and orange "image masks."

Since the registration marks are square and their size relative to each image does not vary much, our code filters contiguous regions of pixels in each image mask by shape and size. The regions that are too large or too small an area to be plausible registration marks are excluded. Likewise, regions that are not approximately square are excluded. The code then sorts the remaining regions from largest to smallest area. For the green image mask, the three largest regions that have approximately equal area are selected as the 3 registration marks. For the orange image mask, the largest region is selected as the registration mark. If there are not enough candidate regions for either green or orange, the code reports this to the user and the coordinates of the fly arena must be input manually.

In Part 2, we use the estimated locations of the four registration marks to collectively transform the images. To start, our code reloads each image, undoing the AGC. This way we can apply AGC again as the final transformation *after* cropping out unimportant details from the images. Then our code rotates each image so that its orange registration mark is in the upper left corner. This ensures all of the images have the same orientation.

Next, our code computes the median location of each of the four registration marks across all images. We do this to reduce noise in the location estimates and to ensure that the transformed images are all the same size. Using the median locations, our code deskews each image so that the outer corners of the registration marks form a rectangle (the fly arena; shown in red in Fig. 1D), then crops each image to this rectangle. Finally, we apply adaptive gamma correction to standardize the brightness of the cropped image, to prepare it for the fly detection phase.

We developed and tested the code for this phase on 15 experimental datasets, composed of 30-50 images each. The code correctly registered the fly arenas and transformed the images automatically for 14 of the 15 datasets. For the 15th dataset, the code reported an error and requested manual input. After manually inputting the coordinates for the fly arena, the code transformed the images.

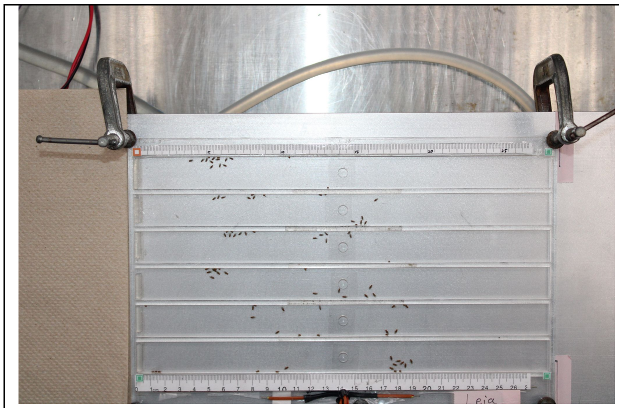

(A)

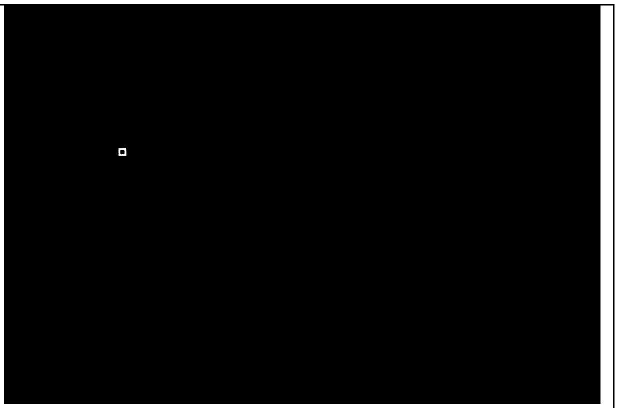

(B)

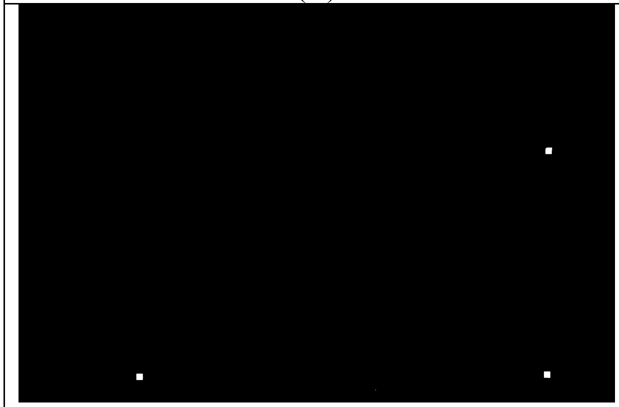

(C)

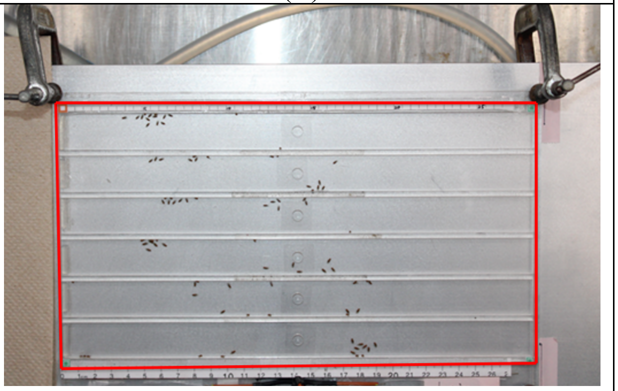

(D)

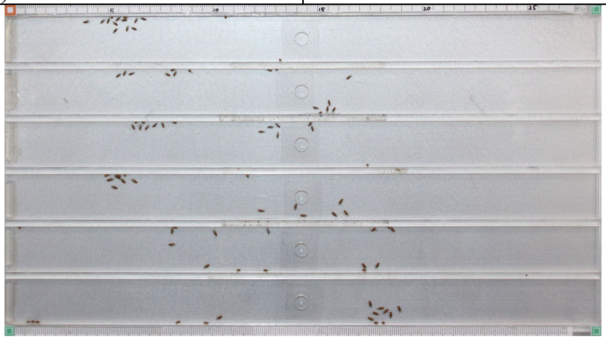

(E)

Figure 1. The transformation of an image in the registration phase: A) the original image; B) the image mask for the orange registration mark, with likely pixels in white and unlikely pixels in black; C) the image mask for the green registration marks, with likely pixels in white and unlikely pixels in black; D) the detected fly arena (red rectangle); E) the deskewed, cropped, and brightness-standardized output image.

#### Phase 2: Detection of Fly Positions for Estimating the Body Temperature

The detection phase locates the flies within each image and estimates their temperatures. To locate the flies, we use an artificial neural network (ANN) called You Only Look Once (YOLO) (Redmon et al., 2016). YOLO is an object detection model that searches for patterns of pixels in order to estimate bounding boxes around specific kinds of objects (in this case, flies; Fig. 2). We estimate the temperature of each fly's location with data from six temperature probes that are placed at equal intervals along the horizontal axis of the fly arena. Each temperature estimate is a linear interpolation based on the fly's predicted bounding box and the two nearest temperature probe measurements (Table 1A ).

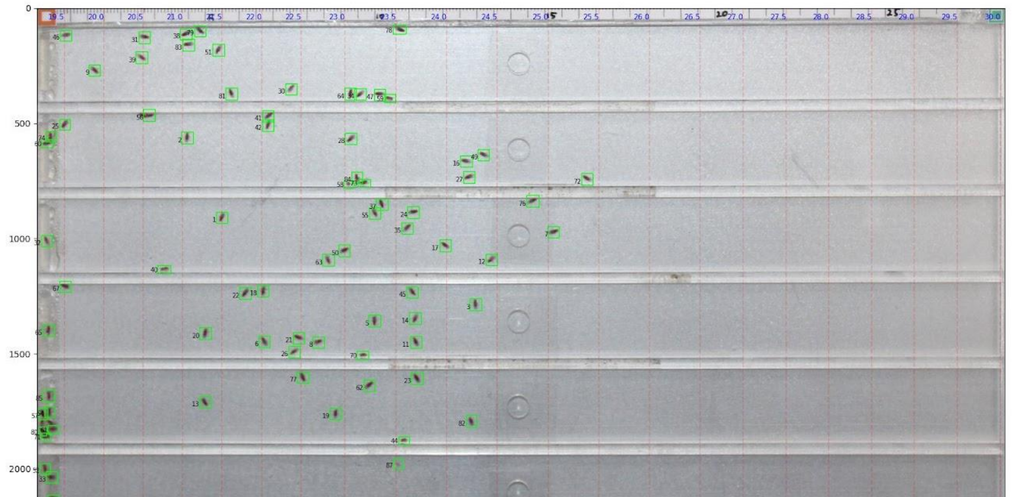


Figure 2. An example of the diagnostic output images from the fly detection tools. The bounding box for each detected fly is shown in green, as well as an identification number. The faint red lines show temperatures, as a fallback for estimating temperatures of flies which were not detected.

#### Model Fine-Tune Training

Training an ANN from scratch requires a large annotated dataset and substantial computing resources. However, the patterns a model learns to solve one problem can generalize to others. As a result, we utilize "transfer learning," a standard practice where an ANN is "pretrained" on a large, general-purpose dataset and distributed for public use (Zhang et al., 2023). The pretrained model can then be "fine-tuned" to specific problems through further training on smaller, problem-specific datasets. We created an annotated fly dataset and used it to fine-tune a pretrained YOLO 8x model (Jocher et al., 2023).

In order to create the annotated fly dataset, we drew bounding boxes around individual flies in X images from 7 experimental datasets. To reduce the amount of labor involved in hand-annotation, we first cropped out a single fly from one of the images and rotated it in 15-degree increments to create a set of 24 rotated templates for matching other flies. We then used code to search each image for regions of pixels relatively similar to the templates and automatically annotate each with a bounding box. We then manually reviewed all images and corrected the bounding boxes, adding them around flies that didn't match the templates (which typically occurred when flies were grouped together or located near the edges of lanes) and removing them from regions that matched but didn't contain a fly (which was rare).

To fine-tune the YOLO 8x model, we used 80% of the images in the annotated fly dataset as training examples (a "training set") and 20% to evaluate model performance during training (a "validation set"). The training procedure for an ANN is iterative, with three steps: 1) estimate the gradient of the loss function on the training set (or a subset); 2) use the gradient to update the model parameters in whichever direction yields the greatest reduction in loss; and 3) evaluate the model on the validation set to assess how well it generalizes to data not in the training set. We repeat these three steps 200 times over the entire training set.

The training procedure evaluates the performance of the model in step 3 by computing the complete intersection over union (CIoU) loss (Zheng et al., 2022) between the predicted bounding boxes and the ground truth (annotated) bounding boxes. CIoU loss is a modification of intersection over union (IoU) loss (Table 3B); IoU loss penalizes predicted bounding boxes for lack of overlap with ground truth bounding boxes. CIoU loss additionally penalizes predicted bounding boxes for center-point distance to ground truth bounding boxes and for aspect ratios that differ from ground truth bounding boxes. Compared to IoU loss, CIoU loss produces greater per-iteration improvement in models during training, and better overall model performance. The model's CIoU loss decreased throughout training on both the training and validation sets (steps 2 and 3; Fig. 3A-B). D decreasing loss on the validation set through the end of training indicates that the model generalizes beyond the training set (it did not "overfit") and may also indicate that it would continue to improve with further fine-tuning. We elected not to pursue further fine tuning because the precision and recall achieved here, which was above 0.9, was sufficient for our purpose.

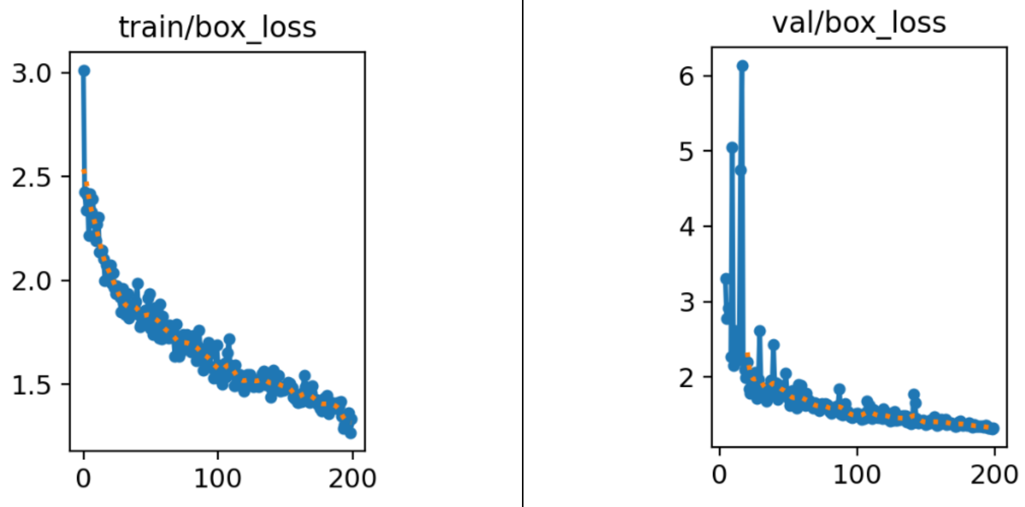


(A) (B)

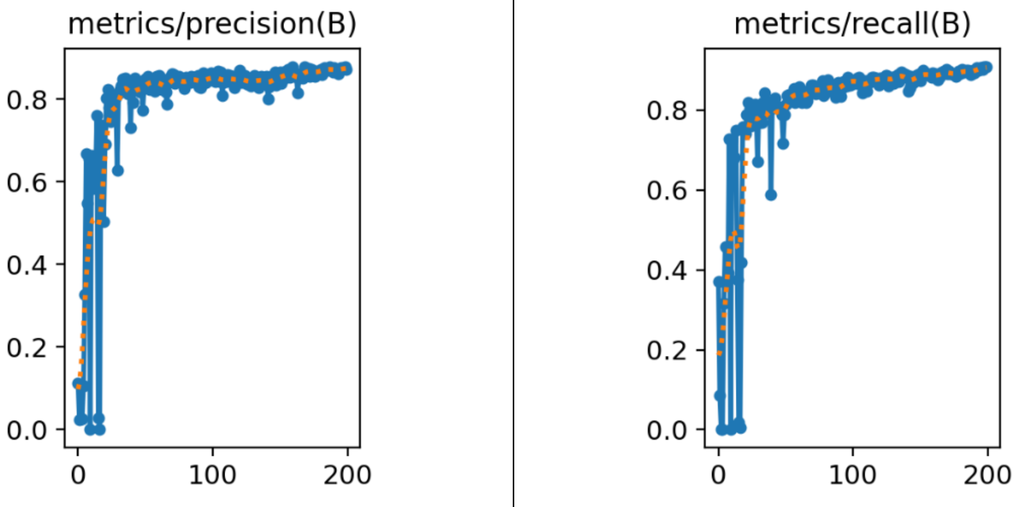


(C) (D)

Figure 3. Plots of summary statistics from fine-tuning the YOLOv8 model: A) loss for bounding box predictions on the training set, B) loss for bounding box predictions on the validation set, C) precision on the validation set, and D) recall on the validation set. In all plots, the x-axis shows the iteration number. Formulas for precision and recall are shown in Table 1C.

To further ensure the model was accurate in its ability to detect flies, we also examined the precision and recall (Table 1C) during training. Precision measures the proportion of predictions that are actually flies. Recall measures the proportion of flies that are actually predicted (that is, how well do the predictions account for all flies). These metrics provide insight into specific kinds of model weakness that are not always apparent from the loss metric alone, such as a tendency to incorrectly predict rare but difficult cases. The precision and recall on the validation set both increased throughout training (Fig. 3C-D), with final precision 0.901 and final recall 0.912.

| |
|---|
| (A) Temperature estimation formula<br>$\hat{T} = (1 - \alpha)T_k + \alpha T_{k+1}$ where $\alpha = \frac{x - x_k}{x_{k+1} - x_k}$<br>for a fly at horizontal location x between temperature probes $k$ and $k + 1$.<br>The variables $x_k$ and $x_{k+1}$ denote the horizontal locations of the probes |
| (B) Intersection over union (IoU) loss |

| |
|---|
| $$IoU\ Loss = 1 - \frac{Area\ of\ Overlap}{Area\ of\ Union}$$ |
| (C) Precision and recall<br>TP : True Positives; TN: True Negatives; FP: False Positives; FN: False Negatives<br>$$Precision = \frac{TP}{TP + FP}$$<br>$$Recall = \frac{TP}{TP + FN}$$ |

Table 1. Formulas related to the temperature estimates and model performance metrics.

#### Phase 3: Cleaning and Post-Processing

After gathering the numerical pixel coordinates and estimated individual body temperatures, we clean the data via Python code. Currently, the assay cannot accurately distinguish whether the flies are in the food region (the first 0.5 cm from the left of the arena) for the food located there or for body temperature (or temperature preference). Because feeding is a confounding variable, all fly data within those regions are automatically cleaned. Separately, we manually clean and remove false positives, yielding an average false positive confidence score of 0.0287 and an SEM of ± 0.0571 (Table 2). The resulting scale of 0.4 and below was taken from the false positive confidence scores to clean false positives stemming from blemishes on the temperature plate. We then use the y-coordinates of each lane in the arena to bin the data by lane. Finally, we average the temperature data points within each lane. We save the binned data points and averages for each lane in CSV files.

| 4-8-24_biden Sample Photo ID # | Average False Positive Confidence Score | Total Sample False Positive N-value | Average Standard Deviation | Average SEM |
|---|---|---|---|---|
| 9906 | 0.01426326 | 5 | 0.003137665 | 0.001403206 |
| 9907 | 0.019028895 | 4 | | |

| | | | | |
|---|---|---|---|---|
| 9908 | 0.022160321 | 10 | 0.005821366 | 0.002910683 |
| 9909 | 0.021539698 | 3 | 0.010719576 | 0.003389828 |
| 9910 | 0.018613597 | 6 | 0.011536251 | 0.006660457 |
| 9911 | 0.010846257 | 1 | 0.005717735 | 0.002334255 |
| 9912 | 0.029121211 | 13 | 0 | 0 |
| 9913 | 0.021357749 | 7 | 0.014347793 | 0.003979362 |
| 9914 | 0.017649492 | 6 | 0.011982601 | 0.004528998 |
| 9915 | 0.024651625 | 7 | 0.009738399 | 0.003975685 |
| 9916 | 0.023500246 | 5 | 0.014152683 | 0.005349211 |
| 9917 | 0.017210707 | 6 | 0.007727283 | 0.003455746 |
| 9918 | 0.02832276 | 7 | 0.008222562 | 0.003356847 |
| 9919 | 0.039971655 | 5 | 0.014334679 | 0.005417999 |
| 9920 | 0.038087603 | 10 | 0.021629774 | 0.009673129 |
| 9921 | 0.038958393 | 8 | 0.029222788 | 0.009241057 |
| 9922 | 0.016808931 | 7 | 0.025070158 | 0.008863639 |
| 9923 | 0.02651968 | 11 | 0.003446561 | 0.001302678 |
| 9924 | 0.018736184 | 9 | 0.011611544 | 0.003501012 |
| 9925 | 0.018611739 | 6 | 0.009576815 | 0.003192272 |
| 9926 | 0.014855462 | 7 | 0.005189814 | 0.002118733 |
| 9927 | 0.039110508 | 10 | 0.003248194 | 0.001227702 |
| 9928 | 0.065175122 | 10 | 0.022700622 | 0.007178567 |
| 9929 | 0.045369094 | 12 | 0.062744251 | 0.019841474 |
| 9930 | 0.024614828 | 7 | 0.040496166 | 0.011690236 |
| 9931 | 0.035881758 | 6 | | |
| 9932 | 0.028075933 | 3 | | |
| 9933 | 0.020666584 | 4 | | |
| 9934 | 0.043873966 | 5 | | |
| 9935 | 0.028788914 | 3 | | |
| 9936 | 0.045841396 | 1 | | |
| 9937 | 0.026103452 | 4 | | |
| 9938 | 0.028285332 | 4 | | |
| 9939 | 0.023412347 | 2 | | |
| 9940 | 0.03665707 | 6 | | |
| 9941 | 0.04110386 | 6 | | |
| 9942 | 0.051488231 | 11 | | |
| 9943 | 0.021467487 | 3 | | |
| 9944 | 0.037919525 | 8 | | |
| 9945 | 0.023687647 | 9 | | |

| | | | | |
|---|---|---|---|---|
| | | | 0.01333777 | 0.005041203 |
| | | | 0.016186171 | 0.006607977 |
| | | | 0.012985692 | 0.007497293 |
| | | | 0.011057598 | 0.005528799 |
| | | | 0.025075088 | 0.01121392 |
| | | | 0.020059525 | 0.011581372 |
| | | | 0 | 0 |
| | | | 0.009364127 | 0.004682064 |
| | | | 0.00668406 | 0.00334203 |
| | | | 0.007331571 | 0.005184203 |
| | | | 0.015114148 | 0.006170325 |
| | | | 0.019651114 | 0.008022534 |
| | | | 0.027686158 | 0.008347691 |
| | | | 0.008839024 | 0.005103213 |
| | | | 0.023054138 | 0.008150869 |
| | | | 0.010275471 | 0.003425157 |
| Total Average | 0.028708463 | 6.425 | 0.0145 | 0.0571 |

Table 2: The average confidence across 40 sampled pictures manually cleaned for false positives.

#### References

Joblove, G. H., & Greenberg, D. (1978). Color spaces for computer graphics. *SIGGRAPH Comput. Graph.*, *12*(3), 20–25. https://doi.org/10.1145/965139.807362

Jocher, G., Chaurasia, A., & Qiu, J. (2023). *Ultralytics YOLOv8* (Version 8.0.0) [Computer software]. https://github.com/ultralytics/ultralytics

Rahman, S., Rahman, M. M., Abdullah-Al-Wadud, M., Al-Quaderi, G. D., & Shoyaib, M. (2016). An adaptive gamma correction for image enhancement. *EURASIP Journal on Image and Video Processing*, *2016*(1), 35. https://doi.org/10.1186/s13640-016-0138-1

Redmon, J., Divvala, S., Girshick, R., & Farhadi, A. (2016). *You Only Look Once: Unified, Real-Time Object Detection*. 779–788. https://www.cv-foundation.org/openaccess/content_cvpr_2016/html/Redmon_You_Only_Look_CVPR_2016_paper.html

Zhang, A., Lipton, Z. C., Li, M., & Smola, A. J. (2023). *Dive into Deep Learning*. Cambridge University Press.

Zheng, Z., Wang, P., Ren, D., Liu, W., Ye, R., Hu, Q., & Zuo, W. (2022). Enhancing Geometric Factors in Model Learning and Inference for Object Detection and Instance Segmentation. *IEEE Transactions on Cybernetics*, *52*(8), 8574–8586. https://doi.org/10.1109/TCYB.2021.3095305
